# Hydroxychloroquine decouples reactogenicity from efficacy in mRNA and viral-vector gene delivery

**DOI:** 10.1101/2025.11.10.687670

**Authors:** Duško Lainšček, Špela Malenšek, Anja Golob-Urbanc, Sara Orehek, Jure Bohinc, Mateja Manček Keber, Timotej Sotošek, Hana Esih, Vida Forstnerič, Peter Pečan, Jelica Pantović-Žalig, Mojca Benčina, Romina Bester, Ulrike Protzer, Roman Jerala

**Affiliations:** Department of Synthetic Biology and Immunology, National Institute of Chemistry, Ljubljana, 1000, Slovenia; Centre for Technologies of Gene and Cell Therapy, 1000 Ljubljana, Slovenia; Graduate School of Biomedicine, University of Ljubljana, Ljubljana, 1000, Slovenia; Institute of Virology, Technical University of Munich, School of Medicine and Health; Institute of Virology, Helmholtz Munich, 81675 Munich, Germany; German Center for Infection Research (DZIF), Munich partner site, 81675 Munich, Germany

## Abstract

Programmable nucleic-acid therapeutics, including lipid nanoparticle (LNP)–formulated mRNA vaccines, genome editors delivered as mRNA, and viral vectors, are transforming precision medicine but remain constrained by innate reactogenicity despite the introduction of chemical modifications into mRNA. Here, we show that mRNA/LNPs and viral vectors elicit a transient inflammatory burst that peaks ∼6 h after dosing in vivo. Co-administration or prophylaxis with hydroxychloroquine (HCQ), a clinically established 4-aminoquinoline, potently attenuated cytokine and chemokine induction across modalities, including strong responses to unmodified mRNA, without compromising therapeutic efficacy. Humoral and cellular immunity to SARS-CoV-2 Spike mRNA vaccines (BNT162b2 and non-modified mRNA/LNP) were preserved, as was live-virus neutralization. Transcriptomic profiling indicated selective dampening of TLR/cGAS–STING pathways with retention of type-I interferon elements compatible with effective vaccination. HCQ further mitigated AAV9-associated blood–brain barrier (BBB) disruption after stereotactic delivery to the mouse brain and reduced mRNA/LNP-associated hepatotoxicity and thrombocytopenia while maintaining therapeutic transgene expression and CRISPR base-editing in vivo. These findings identify HCQ as an anti-reactogenic adjunct that widens the safety window of nucleic-acid therapeutics without sacrificing performance.

---

Nucleic acid (NA)–based therapeutics have evolved from early proposals for gene replacement^1^ to a clinically validated class, spanning prophylactic vaccines^2^, oncology^3^, and genome editing for rare and more common diseases^4^. The COVID-19 pandemic accelerated the adoption of mRNA vaccines delivered by LNPs^5,6^ and revived interest in viral vectors for gene replacement and gene regulation. Despite these gains, an overarching challenge persists: exogenous RNA and DNA are sensed by innate immune receptors as danger signals^7^. Pattern-recognition receptors (PRRs) distributed across endosomes (TLR3/7/8/9)^8,9^ and cytosol (RIG-I/MDA5, cGAS–STING, AIM2, NLRs)^6,10^ recognize several RNA and DNA motifs and trigger inflammatory cytokines, chemokines, and type-I interferons, triggering a strong inflammatory response and side effects.^11–14^ While measured innate activation can be desirable as an adjuvant for vaccines or cancer therapy, excessive or poorly timed responses reduce tolerability, limit dose flexibility, and, for mRNA, directly impede translation and stability^15^.

Sequence engineering, high-purity manufacturing, optimized cap structures, and modified nucleosides such as N1-methylpseudouridine (N1-mΨ) reduce but do not abolish innate sensing^15–17^. Clinical experience reflects this: even chemically modified mRNA/LNPs can cause transient fever, myalgia, and malaise^11,14,18^; unmodified mRNA formulations are even more reactogenic, and viral vectors can trigger systemic or local inflammation, including blood-brain barrier (BBB) perturbation after intracerebral injection^19,20^. These liabilities are particularly constraining when repeated dosing, high exposure, or sensitive tissues (liver, brain) are involved.

Hydroxychloroquine (HCQ), listed among the WHO essential medicines^21^, is a safe and widely-used drug used mainly as an antimalarial agent but also to suppress autoimmunity, mediated by the inhibition of endosomal acidification and masking of nucleic acid accessibility to PRRs and inhibiting their activation^22,23^. These properties suggest a two-pronged mechanism to reduce NA–PRR engagement. We asked whether HCQ could broadly inhibit early reactogenicity without impairing efficacy across mRNA/LNP vaccines, adenoviral vectors, and AAV9, while maintaining antigen expression, immune protection, and genome-editing activity. Here, we show that the prophylactic application and formulation combining NA therapeutics with HCQ strongly reduces the induction of an inflammatory spike. HCQ suppressed the transient inflammation and cytokine signaling by mRNA and adenoviral vaccines without impairing protein translation and neutralization titers. Importantly, we demonstrate that the HCQ pretreatment of experimental animals that underwent stereotactic injection of AAV9 prevented the detrimental BBB damage, supporting the protective properties of HCQ for gene therapy. Strong positive protective effect of the concomitant use of HCQ or as a prophylactic was also found in LNP delivery of therapeutic mRNA or genome editors. LNP-induced liver enzyme elevation, which was the main cause of pausing a certain clinical trial^24^, was significantly reduced in the HCQ-treated animals, showing the potential of this drug.

## A conserved, transient inflammatory burst follows nucleic acid delivery

We first mapped innate immune responses to mRNA varying in cap structure and chemical modification. In human PBMCs electroporated with uncapped, cap-0, or cap-1 transcripts ± N1-mΨ, uncapped and cap-0 mRNA elicited the highest IL-6 and IFNγ; cap-1 N1-mΨ mRNA produced substantially lower but still detectable signals (Fig. 1a–c). Time-course analysis showed cytokines rising by 3 h, peaking at ∼24 h after electroporation, and tapering by 48 h.

**Figure 1.**
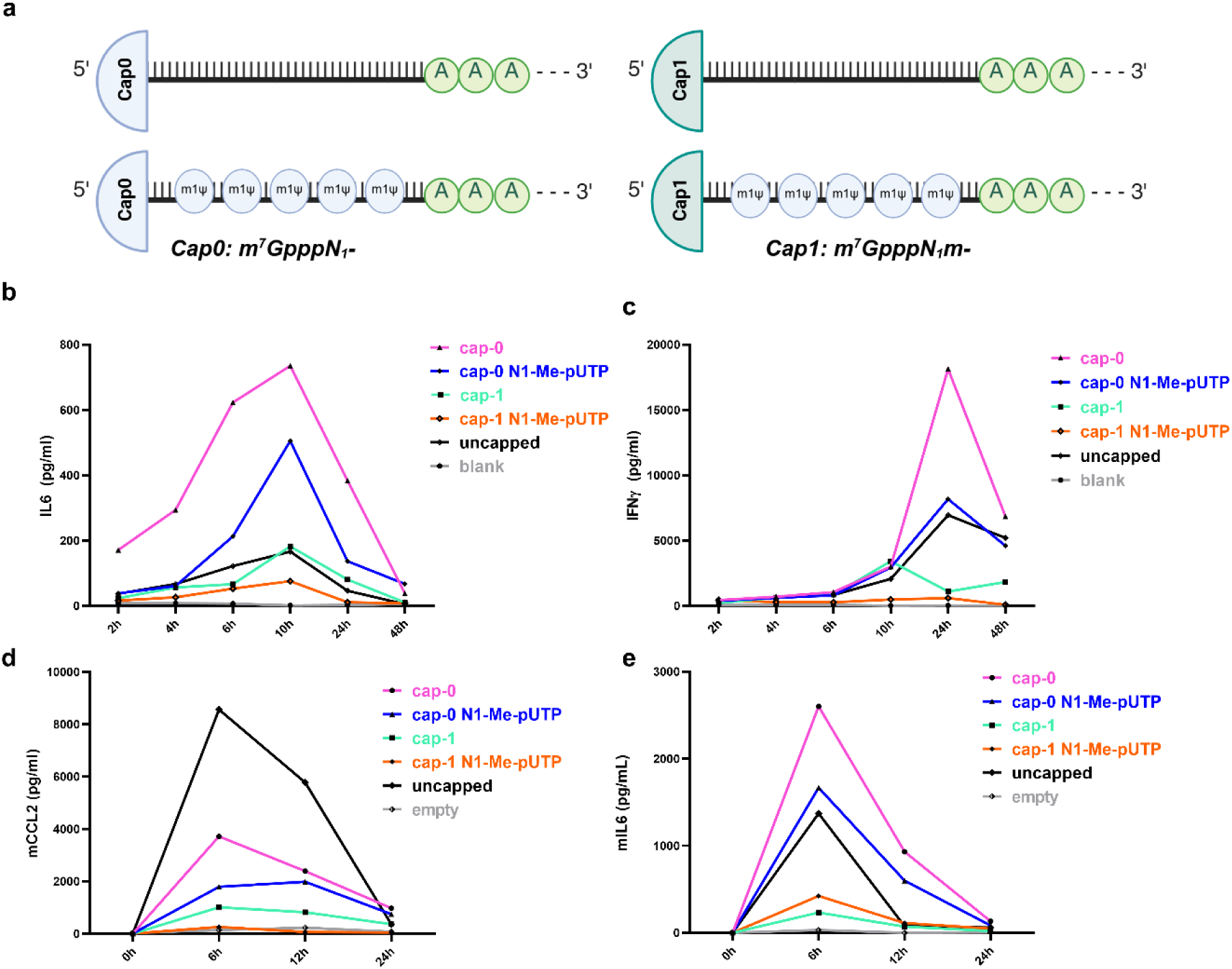
Introduction of mRNA into human cells and mice induces a transient inflammatory spike. a, PBMC electroporation scheme with different mRNA variants (1 µg mRNA per 10 µl tip; luciferase reporter; indicated cap/chemistry). b–c, Secreted human IL6 (b) and IFNγ (c) at 24 h (n=3 donors). d–e, Intramuscular LNP–mRNA (5 µg) in mice produced a cytokine peak at ∼6 h that resolved by 24 h; mouse CCL2 (d) and mouse IL6 (e) (n=3 mice/group). One-way ANOVA with Tukey’s test.

To capture the physiological delivery of NA therapeutics, we formulated the same transcripts in LNPs and injected mice intramuscularly. Serum IL6 and CCL2 surged at ∼6 h and returned toward baseline by 24 h (Fig. 1d-e). As in PBMCs, uncapped and cap-0 mRNAs were most reactogenic, whereas cap-1 N1-mΨ mRNA produced a smaller, yet still substantial spike (Fig.1d-e). The conserved ∼6 h peak suggested a common early innate program triggered across modalities and delivery routes. Vehicle and empty LNPs produced negligible cytokine changes at matched volumes.

## HCQ blunts the inflammation and reactogenicity of mRNA vaccines while preserving the adaptive immune response

We next assessed whether HCQ directly interacts with nucleic acids and LNP vaccine particles. Fluorescence binding assays revealed HCQ association with dsDNA and both ss/dsRNA species, as well as with mRNA/LNP particles (Extended Data Fig. 1a-j). Dynamic light scattering confirmed that adding HCQ to LNP formulations preserved particle size distribution and polydispersity, indicating no disruption of the delivery vehicles (Extended Data Fig. 1k-o).

In PBMCs, HCQ robustly reduced IL6/IFNγ after electroporation of diverse mRNAs. Under the same conditions, bafilomycin A1 (BafA1), an endosomal acidification inhibitor^25^, did not suppress cytokine production, consistent with cytosolic entry bypassing endosomes, supporting the contribution of the direct NA-shielding role for HCQ (Extended Data Fig.2a-e). In mice, co-formulating HCQ with LNP/mRNA strongly suppressed the 6 h cytokine burst (Extended Data Fig.2f-o). Crucially, cap-1 luciferase reporter mRNAs expressed equivalently with or without HCQ in vitro and in vivo. As expected, uncapped transcripts showed low expression due to a rapid decay. Pharmacological congeners (chloroquine, quinacrine) also reduced cytokines in PBMCs (Extended Data Fig.2p-q), suggesting a class effect among 4-aminoquinolines^26^.

We next evaluated clinically relevant vaccines. BNT162b2 (cap-1 N1-mΨ SARS-CoV2 Spike mRNA/LNP) induced the characteristic ∼6 h cytokine peak in PBMCs and in mice; 4-aminoquinolines markedly reduced IL6, IFNγ, and other chemokines in both settings. In PBMCs exposed to BNT162b2, BafA1 also lowered cytokines, which is consistent with the endocytic uptake of LNPs (Extended Data Fig. 3a-d). To test a more stringent condition, we immunized mice with unmodified cap-1 Spike mRNA/LNP, which is more reactogenic than N1-mΨ-containing mRNA. Also, in case of non-modified mRNA, HCQ potently suppressed the cytokine peak at prime and boost (Extended Data Fig. 3e-j) yet did not diminish Spike-specific IgG titres after prime or boost, live-virus neutralization, or CD8⁺ cytotoxicity against Spike-pseudovirus–infected targets (Extended Data Fig. 3k-m). Spectral flow cytometry showed similar frequencies of activated CD4⁺ and CD8⁺ subsets across groups (Extended Data Fig. 3n-o). For the BNT162b2 vaccine, we compared LNP/mRNA-HCQ co-formulation with administration 24 h before vaccination (prophylaxis) in vivo. Both regimens suppressed IL-6, CCL2, and IFN-γ at prime and boost (Fig. 2a–g), with co-formulation providing the most potent suppression. Endpoint Spike-binding titres and live-virus neutralization IC₅₀ were unchanged (Fig. 2h-I, Extended data Fig. 4a). Multiplex panels showed additional broad reductions in eotaxin, G-CSF, IL-5, IL-12p40, KC, MCP-1, MIP-1α/β, and RANTES at 6 h (Fig. 2j, Extended Data Fig. 4b-c) due to the presence of HCQ.

**Figure 2.**
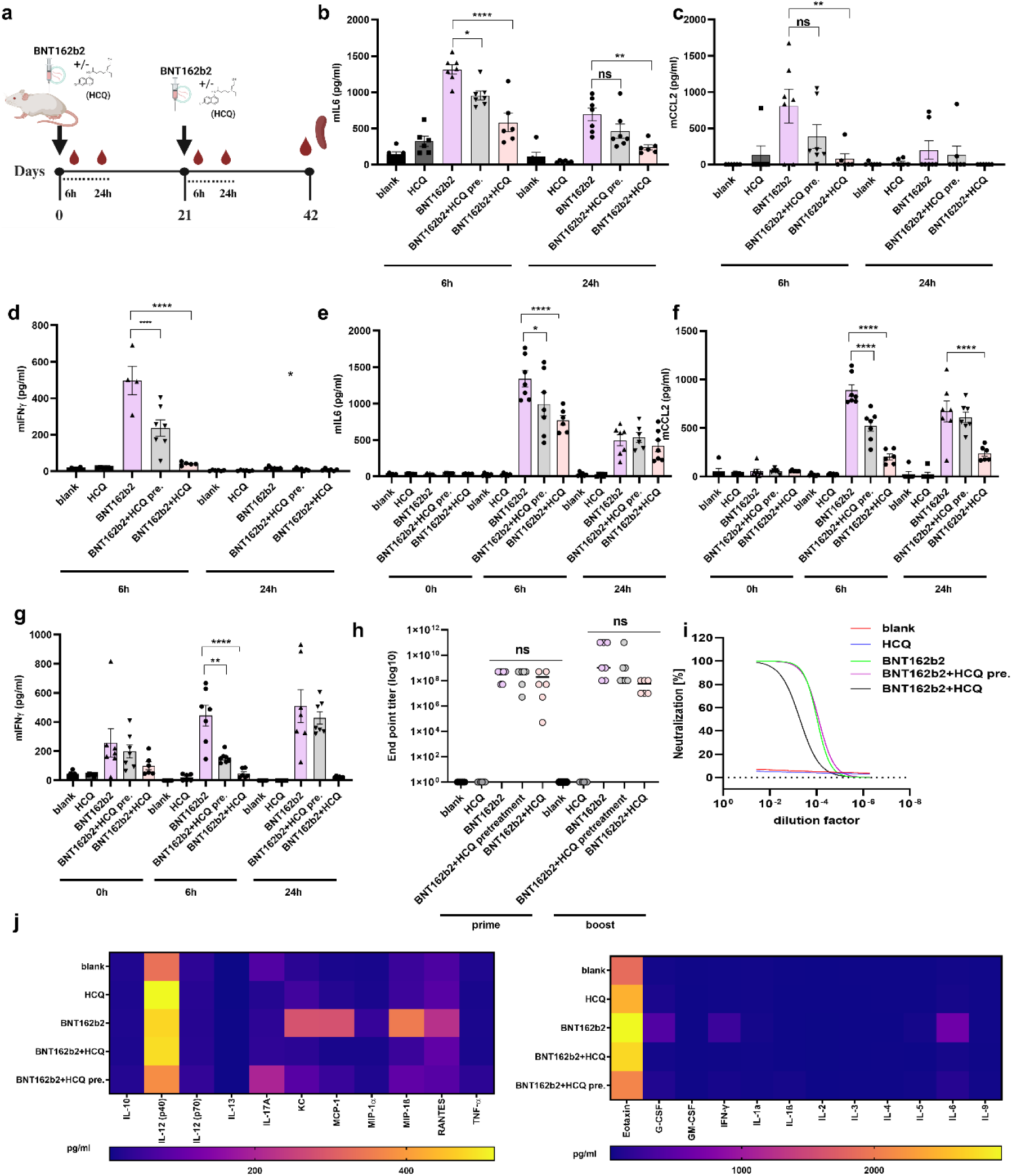
HCQ potently attenuates BNT162b2 reactogenicity without compromising humoral neutralization. **a**, BNT162b2 (4 µg) at days 0 and 21; HCQ co-formulated or given 24 h prior (HCQ pre). **b–g**, Serum IL6, CCL2, IFNγ at 6 h/24 h after prime (**b–d**) and boost (**e–g**). **h,** End-point anti-Spike IgG titres (prime, boost). **i**, Live SARS-CoV-2 neutralization (IC₅₀) across groups. **j**, Multiplex cytokine/chemokine panel at 6 h post immunization in mice (subset shown). n=6; mean±SEM.

CD8⁺ activation and cytotoxicity against Spike-pseudovirus–infected targets remained the same upon HCQ treatment (Fig. 3a-d), whereas cytometry also for the BNT162b2 vaccine exhibits similar frequencies of CD3 cells (Fig. 3e-g, Extended Data Fig. 4d).

**Figure 3.**
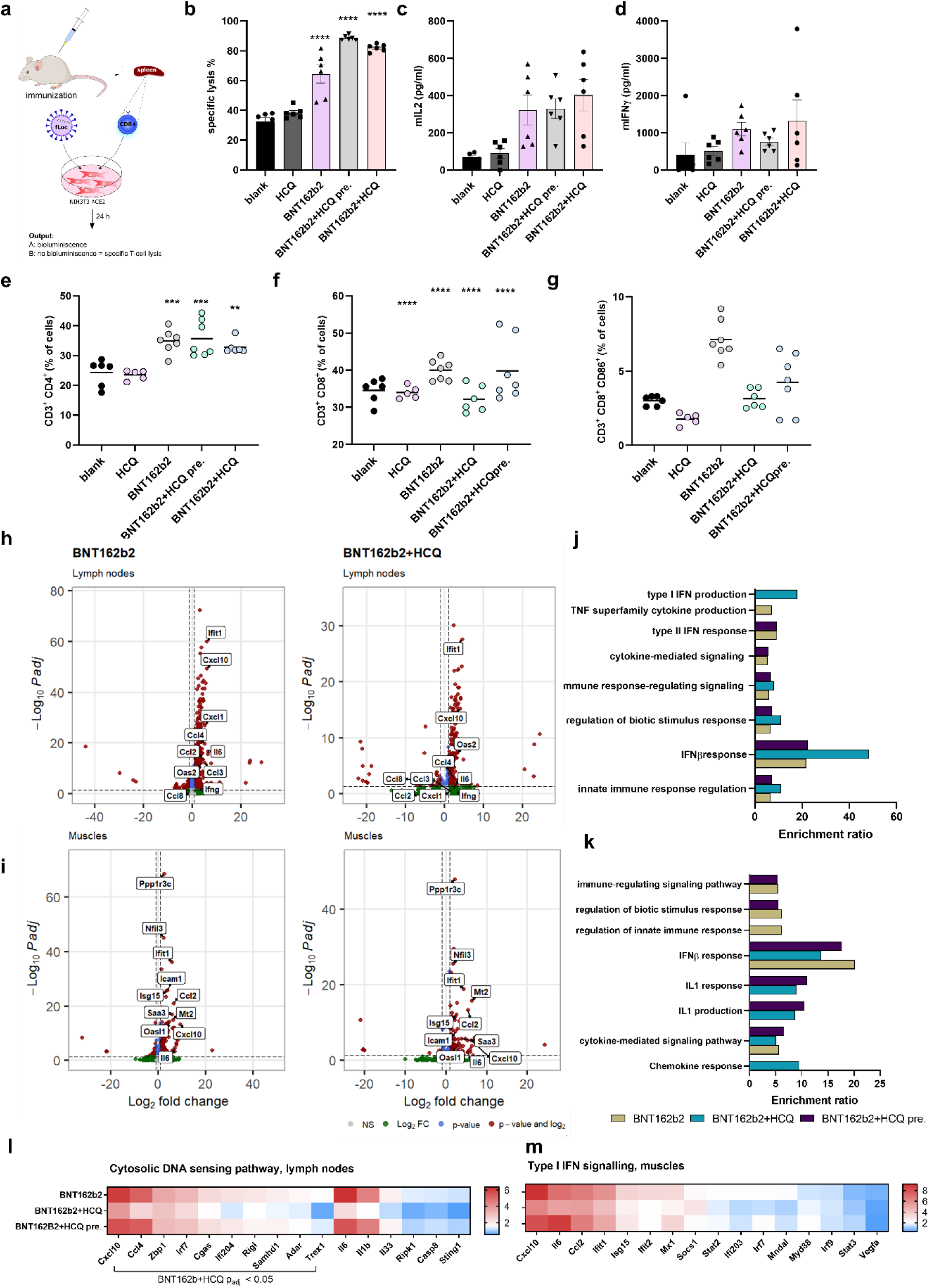
T-cell responses are preserved with innate-signalling programmes selectively dampened by HCQ. **a**, Experimental overview for cellular assays and RNA-seq (3 h post-prime). **b–d**, CD8⁺ specific lysis (**b**) and mouse cytokines IL2 (**c**) and IFNγ (**d**) from co-cultures of BNT162b2 immunized mouse spleen T cells and SARS-pseudovirus infected cells. **e–g**, Spectral flow cytometry of splenocytes (CD4⁺, CD8⁺, CD86⁺CD8⁺) at day 42 post immunization. **h–i**, Enhanced volcano plots highlighting significantly regulated genes in lymph nodes (**h**) and muscle (**i**). **j–k**, Over-representation analysis of upregulated genes. **l-m**, Heatmap of cGAS–STING/TLR pathway genes (lymph nodes). n=6; mean±SEM.

Bulk RNA-seq of draining lymph nodes and peri-injection muscle at 3 h post-prime revealed a distinct innate activation signature in vaccine-only animals (PCA separation by vaccination status), which was selectively dampened by HCQ (Fig. 3h–k; Extended Data Fig. 4e-f). In lymph nodes, HCQ reduced upregulation of TLR/cGAS–STING axis genes and TNF-superfamily cytokine programmes, while preserving a subset of IFN-β-responsive genes potentially supportive of antigen presentation and adaptive priming. In muscle, chemokines and IL-1–responsive transcripts (e.g., Cxcl10, Cxcl2, Cxcl1, Ccl7, Ccl4, Ccl3, Ccl2, Il6) were strongly induced by vaccination; these were blunted by HCQ co-formulation, with prophylaxis producing an intermediate profile (Fig. 3l-m). Clinical chemistry remained unchanged with HCQ (Extended Data Fig. 4g). Together, these results indicate that the pronounced early inflammatory spike is not required for effective priming and that HCQ can temper this spike without impairing antigen expression or adaptive immunity.

## HCQ mitigates adenoviral-vector reactogenicity with preserved efficacy

Because HCQ binds DNA, we examined the chimp-adenoviral vector AZD1222^27^. In splenocyte assays, HCQ suppressed mouse cytokines IL6, CCL2, and IFNγ elicited by vaccine exposure (Extended Data Fig. 5). In vivo, three regimens were tested: co-formulation, contralateral injection (separate site), and 24 h prophylaxis across prime and an 8-week boost. All approaches decreased cytokines at 6 h post-dose (Extended Data Fig. 5a-g). Spike-reactive Ab titers and live-virus neutralization were not reduced by HCQ (Extended Data Fig. 5h-j), and T-cell responses remained intact (Extended Data Fig. 5k-m). Flow cytometry showed no adverse shifts in lymphocyte subsets (Extended Data Fig. 5n-p). DLS confirmed no adverse effects of HCQ on adenoviral particle characteristics (Extended Data Fig. 1l-m).

## HCQ rescues AAV-induced blood-brain barrier damage

Direct intracerebral AAV9 delivery, used frequently to treat neurodevelopmental diseases^28^, to enhance neuronal transduction while limiting systemic exposure, has been linked to blood-brain barrier (BBB) disruption and neuroinflammation^29–31^. We implemented a prophylactic regimen (HCQ daily for 3 d prior, continuing for 2 d after injection) and stereotactically delivered AAV9-eGFP to the mouse cortex (Fig. 4a). Mouse IL6 was reduced at days 1 and 30 with HCQ in comparison to the AAV9 alone, and in vivo eGFP signal indicated preserved transduction efficiency (Fig. 4b; Extended Data Fig. 6c).

**Figure 4.**
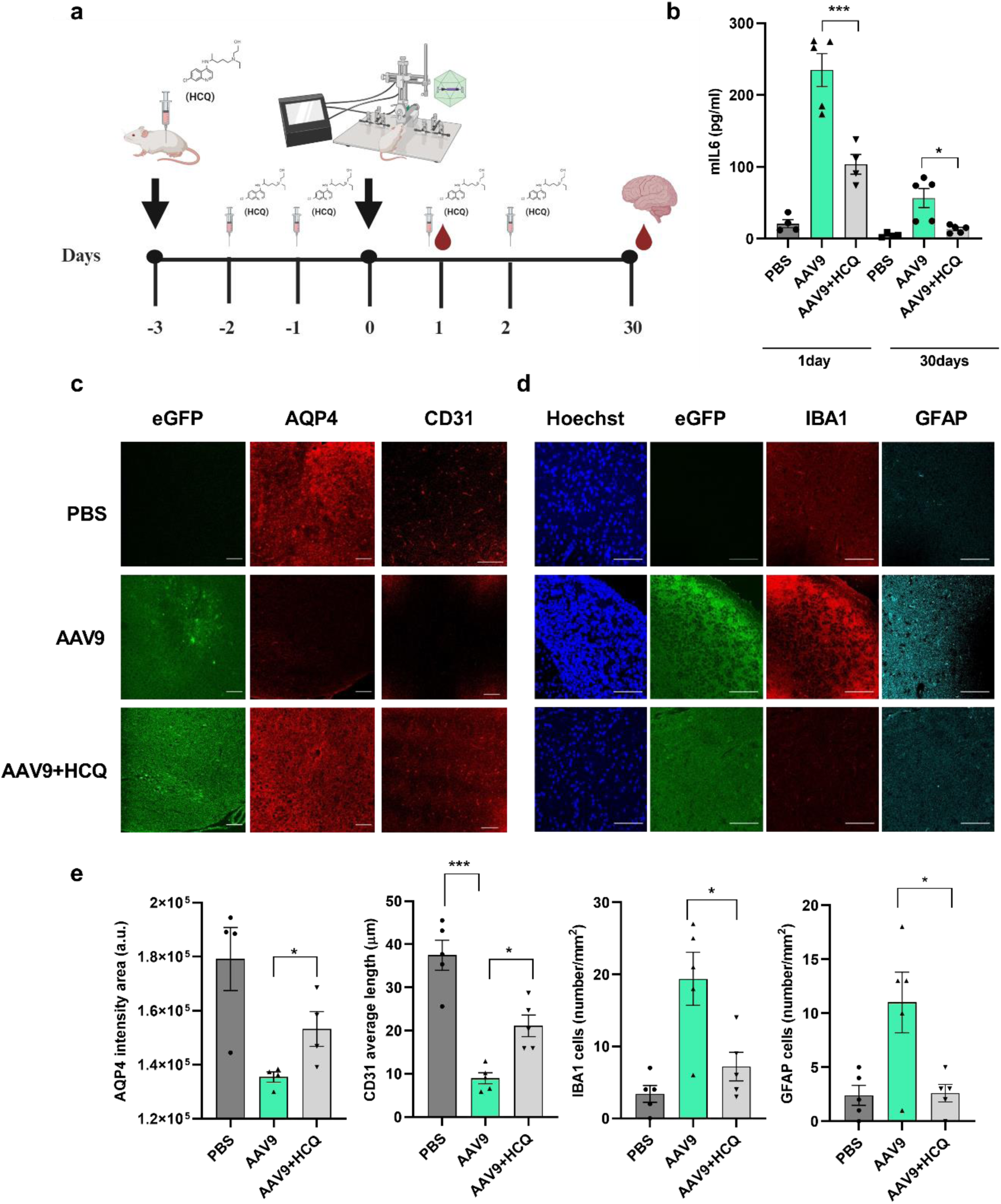
HCQ reduces blood–brain barrier damage after stereotactic AAV9 injection. **a**, HCQ (60 mg kg⁻¹) for 3 d before AAV9-eGFP stereotactic injection (1×10¹⁰ pfu, cortex) and for 2 d after. **b**, Serum mouse IL6 at days 1 and 30 post injection (n=5; mean±SEM). **c–e**, IHC quantification at day 30 post injection: AQP4 and CD31; IBA1 and GFAP; representative images; scale bars 100 µm.

Immunohistochemistry at day 30 revealed BBB injury in AAV9-only brains: reduced CD31 (endothelial) length, diminished AQP4 perivascular staining, increased IBA1⁺ microglia and GFAP⁺ astrocytes, and lower NeuN (neuronal nuclei) (Fig. 4c-e). HCQ prophylaxis markedly preserved CD31 and AQP4 and reduced microgliosis and astrocytosis (Fig. 4e), with higher NeuN abundance (Extended Data Fig. 6d). Thus, HCQ safeguards vascular and glial homeostasis during intracerebral vector administration, rescuing BBB damage without compromising transduction, thus therapeutic efficiency.

## HCQ ameliorates LNP-mRNA hepatotoxicity and thrombocytopenia and supports base-editor dosing

LNP/mRNA exposure often concentrates in the liver, where dose-dependent hepatotoxicity^32–34^ and thrombocytopenia have been observed clinically and preclinically^24^, which prevents repeated treatment that could be used to deliver therapeutic mRNA to generate the desired proteins. We first confirmed therapeutic expression using cap-1 Epo mRNA/LNPs that could be used to treat anemia^35^. Serum EPO rose and hematocrit increased appropriately, with HCQ leaving both readouts unchanged. Importantly, however, IL6 and CCL2 at 6 h were strongly suppressed by HCQ (Fig. 5a-d). Dose-escalation with eGFP mRNA/LNPs (0.2–1.5 mg kg⁻¹) revealed 6 h cytokine peaks across doses; at 1–1.5 mg kg⁻¹, ALT/AST rose significantly by day 4, whereas empty LNPs did not (Extended Data Fig. 7), confirming that the nucleic acid cargo was the main driver of inflammation in a dose-dependent manner. HCQ co-administration blunted cytokines and reduced ALT/AST elevations across reporters and Epo (Fig. 5e–f; Extended Data Fig. 7i), with liver histology showing diminished inflammatory foci and hepatocyte injury (Fig. 5g; Extended Data Fig. 7l). Complete blood counts showed that HCQ mitigated LNP-mRNA–induced thrombocytopenia while preserving EPO-driven erythropoiesis (Extended Data Fig. 8).

**Figure 5.**
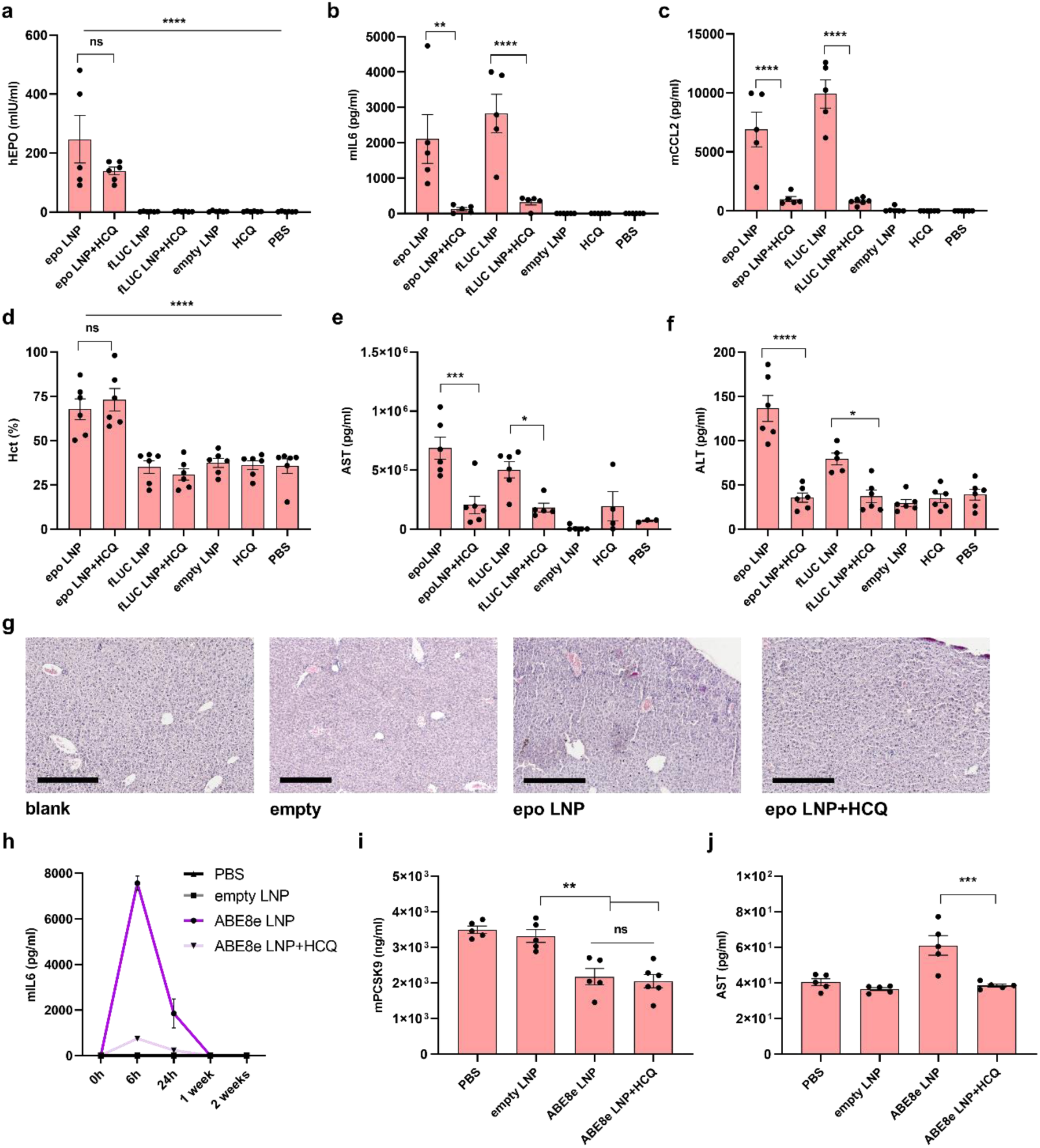
HCQ mitigates LNP–mRNA hepatotoxicity and thrombocytopenia and increases the safety of multiple supplementary gene dosages and in vivo base-editing. **a-c**, Erythropoietin (Epo) or fLUC mRNA/LNP (1 mg/kg of body weight) ± HCQ (60mg of body weight) every 72 h for 1 week. Human Epo **(a)**, mouse IL6 (**b**) and CCL2 (**c**) at 6 h after first dose. **d**, Hematocrit values at day 8. **e–f**, AST and ALT liver enzymes at day 4. **g**, Representative liver H&E (day 8); scale 300 µm. **h–j**, ABE8e + *Pcsk9* gRNA mRNA/LNP (1.5 mg/kg of body weight) ± HCQ (60 mg/kg body weight). Re-dosage was performed three days later. Mouse IL6 at 6 h post first injection (**h**), serum mouse PCSK9 at day 21 (**i**) and AST at day 4 (**j**); mean±SEM; n=5–6.

A prominent emerging application of mRNA is to deliver genome editors, to repair or inactivate the target genes^36,37^, such as e.g., *Pcsk9*, whose inactivation provides strong protection against cardiovascular diseases^32^. Base editing of a splicing site of *Pcsk9* represents a highly promising approach to treat one of the most common causes of mortality, as it enables lasting silencing without cleavage of the DNA. The key obstacle to its clinical development is linked to the hepatotoxicity of the delivered LNP/base editor mRNA. To examine the effect of HCQ on the suppression of inflammatory effects, we delivered ABE8e base editor mRNA with *Pcsk9* gRNA in LNPs at a therapeutically relevant dose. HCQ reduced early cytokines (Fig. 5h) and lowered AST/ALT elevations (Fig. 5j; Extended Data Fig. 9) while the LNP/mRNA caused a desired decrease of serum PCSK9, resulting from genome editing of the *Pcsk9* gene (Fig. 5i). These data support HCQ as a generally applicable adjunct to improve tolerability of repeated or higher-dose LNP-mRNA regimens, including genome editors.

## Discussion

This study identifies a conserved early innate response, peaking approximately six hours after dosing, that follows nucleic-acid delivery across LNP-mRNA, adenoviral, and AAV9 platforms. We show that hydroxychloroquine (HCQ) consistently decouples this transient reactogenicity from functionality: antibody titers, live-virus neutralization, CD8⁺ cytotoxicity, transgene expression, and genome editing are all preserved, while blood–brain barrier disruption, liver enzyme elevations, and thrombocytopenia are mitigated.

Even when therapeutic nucleic acids are chemically identical to the cellular nucleic acids^16,38^, they may trigger an innate immune response through aberrant localization (cytosolic DNA, endosomal RNA or DNA). Before associating with RNA-binding proteins, such species may activate pattern-recognition receptors and trigger inflammatory cascades^6,39^, leading to the transient cytokine response that precedes translation.

Our findings support a multifactorial protective role for HCQ. First, direct nucleic acid binding likely limits receptor engagement: HCQ associated with dsDNA and RNA species and reduced cytokine induction even after electroporation, where endocytosis is bypassed. Second, endosomal pH modulation by HCQ can interfere with TLR ligand generation via RNAse T2 processing^17^ and inhibit the proteolytic maturation of TLR7/8/9^40^, consistent with the BafA1 sensitivity observed for BNT162b2-induced stimulation in PBMCs. Transcriptomic profiling revealed attenuation of TLR/cGAS–STING pathways while preserving IFN-β response elements, molecularly reconciling reduced inflammation with maintained vaccine priming.

Consistent with previous reports^40^, we confirm that the mRNA payload is the principal driver of LNP/mRNA reactogenicity, whereas the lipid components alone are not sufficient at the doses tested, in agreement with the tolerability of empty LNPs. In the liver, HCQ lowered ALT/AST levels and rescued platelets, supporting its potential in repeated dosing or high-exposure regimens such as chronic protein replacement or in vivo gene editing. In the brain, where inflammation and barrier disruption are particularly detrimental, HCQ reduced microgliosis and astrocytosis and preserved endothelial and perivascular integrity following AAV9 administration, suggesting a route to safer intraparenchymal gene delivery.

HCQ is inexpensive, orally bioavailable, and clinically familiar. Co-formulation, where it associates with the LNP cargo without altering particle size or encapsulation efficiency or temporally coordinated dosing, may provide a practical means to improve tolerability without broad immunosuppression, distinguishing this approach from glucocorticoid co-therapy^41^. While optimization of dosing, timing, and formulation remains to be defined, HCQ and related 4-aminoquinolines emerge as anti-reactogenic adjuncts capable of broadening the therapeutic window of nucleic-acid-based medicines.

Our experiments were performed in mice and ex vivo human cells. Translation to humans will require pharmacokinetic and exposure matching, particularly for CNS delivery. We did not assess chronic or multidose regimens beyond the tested schedules. Although HCQ did not impair vaccine immunogenicity, applications requiring strong adjuvanticity, such as cancer vaccines, may not benefit. Finally, because PRR engagement differs across nucleic acid sequences, chemistries, and formulations, the effects of HCQ should be validated case-by-case.

## Methods

### Cells

PBMC cells for nucleic acid stimulation and electroporation were obtained from healthy donors. Samples were obtained with informed consent, and according to the study protocol approved by the National Medical Ethics Committee (0120-199/2022/3) for conducting research on determining the immune response of COVID-19 vaccines. The cells were maintained in RPMI1640 (Invitrogen Life Technologies) cell media, supplemented with 10% (v/v) heat-inactivated FBS (Invitrogen Life Technologies).

Mice splenocytes for nucleic acid stimulation were obtained from mice spleens of different strains of mice, according to the directive of the EU 2010/63. Tissue collection was approved by the Administration of the Republic of Slovenia for Food Safety, Veterinary Sector and Plant Protection of the Ministry of Agriculture, Forestry and Foods, Republic of Slovenia (Permit Number U34401-13/2022/3). Single cell suspensions from spleens without enzymatic treatment were obtained using the tissue dissociator gentleMACSTM Dissociator, according to the manufacturer’s instructions (MiltenyiBiotec, Bergisch Gladbach, Germany). Cells were maintained in RPMI1640 cell media, supplemented with 10% (v/v) heat-inactivated FBS. The cells were treated with various nucleic acid-based vaccines, combined with HCQ. To determine cell death LDH cytotoxicity assay (ThermoFisher Scientific) was carried out according to the manufacturer’s protocol.

### Cell electroporation

PBMCs (2×10^7^ cells/ml) were electroporated by Neon electroporation system (ThermoFisher Scientific), using T buffer in 10 μl electroporation tips (electroporation parameters: 2150V voltage, 20 ms pulse width, 1 pulse). The total amount of mRNA per electroporation was 1 μg.

### Lipids and hydroxychloroquine

Ionizable cationic aminolipid heptatriacont-6,9,28,31-tetraene-19-yl 4-(dimethylamino) butanoate (DLin-MC3-DMA; MC3, HY-112251), hydroxychloroquine sulfate (HCQ; HY-B1370) and quinacrine dihydrochloride (Q) were purchased at MedChemExpress. 1,2-diastearoyl-sn-glycero-3-phosphocholine (DSPC; SI-850365P) and cholesterol (Chol; C8667) were purchased at Sigma. 1,2-dimyristoyl-glycero-3-methoxypolyethylene glycol-2000 (DMG-PEG2000; 880151P) was purchased at Avanti lipids. Chloroquine sulfate (CQ) was purchased from Sigma.

### IVT mRNA synthesis

The mRNAs were prepared by in vitro transcription (IVT) with T7 RNA polymerase according to the manufacturer’s instructions (Ribomax LargeScale RNA production System, Promega P1300) or stated otherwise. We used purified PCR fragments for DNA templates, with already encoded 102bp long polyA tail. dsRNA was synthesized using DNA templates with T7 promoters on both 3’ and 5’ ends. For integration of modified uracil nucleotides, Pseudo-UTP (Jena Bioscience NU-1139S) and N-methylpseudo-UTP (Jena Bioscience NU890-S) we added 20uL of each, replacing non-modified UTP in the reaction mixture entirely. After DNase I treatment of the IVT reaction mix, we performed capping of mRNA, using Vaccinia capping system (NEB M2080S) for formation of Cap-0 and additionally, Cap 2-O’methyltransferase for formation of Cap-1 (NEB, NE-M0366S). mRNA was purified according to the manufacturer’s instructions (Phenol-Chloroform extraction) or with NEB’s Monarch RNA Cleanup Kit NEB T2040L. mRNA’s integrity and correct size were assessed using Agilent’s 2100 Bioanalyzer with Agilent RNA 6000 Pico kit (5067-1513, 5067-1514), concentration and A260/A280 ratios were measured on Nanodrop.

### LNP formulation and characterization

Lipid nanoparticles, encapsulating mRNA, were prepared as previously reported ^49^, using Precision Nanosystems Ignite microfluidic mixing platform. Briefly, we first prepared a lipid formulation in ethanol at molar ratios of 50.5:10:38:1.5 (MC3/DSPC/Cholesterol/DMG-PEG2000). Thus, a prepared lipid mix with a final lipid concentration of 12.5mM and mRNA concentration of 0.174-0.200 mg/mL in 25mM citrate buffer (pH 4.0) was injected into the NxGen cartridge at flow rate ratios 1:3 with a final flow rate of 12mL/min. N/P ratio was 4 (cationic ionizable lipid to anionic mRNA). Subsequently, we diluted the product 10 times in phosphate buffer saline (pH 7.4) and concentrated the diluted product at 2000g for 30min. LNPs were filtered through 0.22uM membranes and stored at 4°C or at −80°C (diluted 2x with 10% sucrose solution). BNT162b2 mRNA/LNP and chimp-adenoviral vector AZD1222 doses against the SARS-CoV-2 ancestral strain that remained unused were used, based on the approval of the Medical Ethics Board of Slovenia and local authorities at Regierung Oberbayern, Munich, Germany.

In-house made LNP’s and BNT162b2 vaccine’s size and distribution were characterized with dynamic light scattering (measured on Malvern Zetasizer). mRNA encapsulation efficiency and concentration inside LNPs was determined using QuantiTRibogreen RNA assay (Molecular Probes R-1141) and Quantifluor RNA system (Promega E3310).

### Fluorescence spectroscopy

HCQ (1 mM) was mixed with various concentrations of nucleic acids, LNPs or vaccines (AZD1222 and BNT162b2) at a room temperature in 96 well black microplate. In the competition assay the PI (50 μM) was added to the mixture. Fluorescence measurements were performed with a microplate reader (BioTek) using excitation at 330 nm for HCQ and excitation at 535 nm for PI. The emission spectra of HCQ and PI was recorded between 450 and 580 nm or 550 and 750 nm, respectively. All compounds were dissolved in miliQ.

### RNA sequencing

Total RNA was extracted from homogenized lymph node and muscle tissue with TriPure^TM^ Isolation Reagent (Merck) and High Pure RNA Isolation Kit (Roche) following manufacturers’ instructions. Purity (ratios A260/280 and A260/230) and quantity were measured on Nanodrop spectrometer (Thermo Fisher Scientific). Three biological repeats of each sample were sent for sequencing at Azenta Life Sciences (Genewiz, Leipzig). They performed sample quality control, library preparation, rRNA removal by polyA selection for mRNA species from muscle tissue and rRNA depletion for lymph nodes due to lower quality of the RNA samples. The libraries were run on Illumina® NovaSeq^TM^, using 2×150bp paired-end sequencing protocol, with sequencing depth of 20-30 million reads per sample. First, we performed QC with FastQCheck and then paired-end adapter trimming, using CutAdapt^43^. Then we “pseudo-aligned” trimmed samples to mouse genome, using Kallisto pseudo alignment^44^ with pre-produced index and gtf files, downloaded from Kallisto transcriptome indices data source (version June 22 2019; kallisto 0.45.1, Ensembl v96 transcriptomes). Differential expression was performed using Rstudio (RStudio 4.3.2 (2023-10-31 ucrt)) with the R-package DESeq2^45^. Data was normalized with the built-in function median-of-ratios, threshold of significance was set to adjusted P-value < 0.05. Volcano plots were drawn with EnhancedVolcano^46^ Bioconductor package. To perform enrichment analysis, we took significantly upregulated genes of each sample (adj. P < 0.05, log_2_FC > 0) as an input to WEB-based Gene Set Analysis toolkit (WebGestalt)^47^. We performed ORA (over-representation analysis) and for annotation we chose the functional database gene ontology (non-redundant biological process). The data was submitted to the GEO repository (GSE277440).

### ELISA assays

Values of human IL6 and IFNγ were determined by using Human IL2 ELISA Kit (Invitrogen) and Human IFNγ (eBioscience) according to the manufacturer’s protocol. Mouse IL6, CCL2, IL2, and IFNγ were determined by IFN gamma ELISPOT ELISA Ready-SET-Go!™ Kit; (15531137, eBioscience, San Diego, CA, USA) according to the manufacturer’s protocol. Human EPO values were determined by using the Human EPO ELISA kit (ThermoFisher Scientific) following the manufacturer’s instructions.

### Bio-plex analysis

Multiplexing of mouse cytokines and chemokines was determined by using Bio-Plex Pro Mouse Cytokine Grp I panel 23-plex (Bio-Rad) according to the manufacturer’s instructions. For readouts Bio-Plex 200 System was used with Bio-Plex Manager, version 6.2 (Bio-Rad). **Spectral Flow cytometry analysis**

Cells (4-5 × 10^6^) were resuspended in FACS buffer (150 µl, PBS supplemented with 10% FBS). For Live/Dead staining, ZombieNIR dye was used according to the manufacturer’s protocol. Briefly, the FACS buffer was replaced with 100 µl PBS containing ZombieNIR dye (dilution 2000 to 5000). Samples were incubated for 10-15 min on ice, and the reaction was stopped with the addition of FACS buffer (100 µl). The ZombieNIR dye was removed with centrifugation at 450 rpm, and the collected cells were resuspended in 50 µl FACS buffer containing 2 µl Mouse TrueSatin FcX (BioLegend) or anti-mouse CD16/CD32 (Fcy III/II receptor) (BD Pharmingen). Ten minutes later, an antibody cocktail was added (50 µl) to the mixture, and samples were incubated for at least 30 minutes on ice. The antibody cocktail was prepared in FACS buffer containing 5 µl True Stain Monocyte blocker (426103, BioLegend). See Tables 1 and 2 for a list of the antibodies used in this study. Before analysis, cells were washed with FACS buffer, and fluorescence was measured on the 3-laser Aurora spectral flow cytometer (Cytek Bioscience). All antibodies were titrated individually according to standard practice before being used in the panel. As reference controls, an unstained sample and, for every color, a single-stain reference control were acquired. All reference controls underwent the same protocol as fully stained samples, including washes. Reference controls were acquired once and used for unmixing the multiple batches. For all parameters, a cell type of interest was used. Data were acquired and unmixed using SpectroFlo v3.1.0 software (Cytek Bioscience) using the same instrument settings for every run. The resulting unmixed FCS files were analyzed using manual gating in Flowood v software (BD Biosciences). First, a manual data check was performed to ensure the exclusion of technical artefacts and bad-quality samples (clogs, doublets, and dead cells). The unmixing of raw data was performed using single-stain controls as references (no manual compensation was used). FlowAI software was used to exclude noise, anomalies in flow rate, signal acquisition, and outlier events (REF). Next, doublets and dead cells were removed, and CD45-positive cells were gated manually, subsampled using the FlowJo_Downsample function and saved as new FCS 3.1 files. Before dimensionality reduction and clustering the new files were merged using the FlowJo_Concatenation function. The manually gated cells were further analyzed using the UMAP dimensionality reduction method and X-shift/FlowSOM clustering approach. For T cells dimensionality reduction: CD4, CD8, MHCII, CD103, CD11b, CD86, CD69 and CD62L* and CD44*, markers were used; for B cells: MHCII, CD62L*, CD44*, CD86, CD161, CD11c and for the rest of cells: CD103, CD44*, CD4, CD62L*, CD11c, MHCII, CD11b, CD8a, CD64, Ly6C, CD86, CD69. FCS files are available upon request.

### Mouse studies

All animal experiments were performed according to the directives of the EU 2010/63 and were approved by the Administration of the Republic of Slovenia for Food Safety, Veterinary Sector and Plant Protection of the Ministry of Agriculture, Forestry and Foods, Republic of Slovenia (Permit Numbers U34401-9/2022/4; U34401-6/2025/8; U34401-11/2025/8). Laboratory animals were housed in IVC cages GM500 (Techniplast), fed standard chow (Mucedola) and tap water were provided ad libitum. The cages were enriched using Nestlets nesting material and mouse houses. Mice were maintained in a 12-12-hour dark-light cycle at approximately 40-60% relative humidity with 22°C of ambient temperature. All animals used in the study were healthy, accompanied by a health certificate from the animal vendor. Health/microbiological status was confirmed by the FELASA-recommended Mouse Vivum immunocompetent panel (QM Diagnostics).

To test the effect of HCQ on nucleic acid immunizations, female 8-10 weeks C57Bl6/J OlaHsd mice were used. Mice were intramuscularly immunized with Covid19 vaccine (BNT162b2 (Pfizer); 4μg/animal, AZD1222 (AstraZeneca); 3,5×10^4^ pfu/animal or in house prepared LNPs, containing Spike mRNA); 4μg/animal, expressing Spike protein of interest, combined with hydroxychloroquine sulfate (60mg/kg of body weight of the animal), combined into a single injectable composition if not stated otherwise. COVID-19 vaccines were technical waste, obtained from Golnik hospital or from TUM. In some cases, HCQ was administered 24h before the vaccine. Immunization was carried out under general inhalation 1,8% MAK isoflurane anesthesia (Harvard Apparatus). Vaccine (prime; 1^st^ dose) was administered using a 30G needle into *m. tibialis anterior* after appropriate area preparation. The boost (2^nd^ dose) was administered 3 weeks later. The blood was drawn 6 and 24 hours after prime. The blood was taken again before the 1^st^ boost and again 6 and 24 hours after the 1^st^ boost. Blood was drawn from the lateral tail vein using Microvette 300 (Sarstedt). Three weeks after the boost, the experiment was terminated. Final blood was taken, and spleens were harvested from animals. Mouse sera were prepared by centrifugation of blood samples at 3000RPM/ 20min at 4°C.

To study the effect of LNP-induced liver damage and thrombocytopenia, a mixture of HCQ (60mg/kg) and LNPs (0,08mg/kg or 0,5 mg/kg of body weight) containing in vitro transcribed human EPO mRNA or fLUC mRNA or eGFP (were injected intravenously in C57Bl/6J OlaHsd (female mice, aged 8-10 weeks) every 72 h for one week. Blood was taken at 6 hours post-first injection and then at day 8.

Complete biochemical profile was determined using VetScan Mammalian Liver Profile reagent and Comprehensive Metabolic Panel rotor and analyzed on the biochemistry analyzer VetScan VS2 (Abaxis). Five-part complete blood count was performed using Vetscan^®^HM5 Hematology Analyzer (Abaxis).

### Mouse *Pcsk9* in vivo base editing

LNPs, containing ABE8e mRNA and gRNA (CCCATACCTTGGAGCAACGG^32^, purchased from Synthego), which was in vitro transcribed using mMESSAGE mMACHINE™ T7 mRNA Kit with CleanCap™ Reagent AG (ThermoFisher Scientific) were injected into Balb/c OlaHsd female mice, aged 6 weeks at a dose of 1,5 mg/kg of body weight, followed by another dose 2 days later. Blood was taken to evaluate cytokines, liver enzymes, and mouse PSCK9 level using ELISA (Sinobiological). Three weeks later, organs were harvested and pathohistological analysis was performed. After 48 h of fixation in 10% neutral buffered formalin (Sigma-Aldrich) and gradient ethanol dehydration, the tumor samples were embedded in paraffin (Leica Biosystems), sectioned into 7 μm pieces using Microtome (Leica Biosystems) and mounted on adhesive-coated slides (Leica Microsystems). After deparaffinization and hydration, sections were stained with Mayer’s Hematoxylin Solution (Merck) and Eosin Y-solution (Sigma-Aldrich). Histology slides were scanned using Ocus®40 (Grundium), whereas pictures were analyzed using Aperio ImageScope v12.4.6.5003 (Leica).

### Inducing blood-brain barrier damage

To induce blood-brain barrier damage, female 10 weeks old C57Bl6/J OlaHsd mice were used. By stereotactic injection, described as in^20^ AAV9-GFP (Origene) was injected into the right cortex (AP +0.6 mm, ML −2 mm, DV −0.7 mm) by using 71000 Automated Stereotaxic Instrument (RWD). Each animal was injected with 1×10^10^pfu of AAV viruses in a total volume of 1,5μl. The next day and on day 30, the blood was taken to check for the mouse IL6 values. On day 30, eGFP fluorescence was determined by using the spectral unmixing mode in vivo imaging system using IVISIII (Perkin Elmer, Waltham, MA, USA). Fluorescence values are presented as average radiant efficiency (p/s/cm2/sr; μW/cm^2^), which were determined using

Living Image® software. The animals were humanely sacrificed on day 30, the brains were removed, snap frozen in liquid nitrogen and used for further immunohistochemistry analysis. Using a cryostat MEV (SLEE medical GmbH), 6 μm brain tissue slices were cut. Cell nuclei were stained with Hoechst dye (Invitrogen). For GFAP detection, mouse GFAP (GA5) (Cell Signaling Technology; dilution 1:100) antibody was used. Alexa Fluor 568 goat anti-mouse IgG (H+L) 2mg/ml (Invitrogen; dilution 1:1000) was used as a secondary antibody. For CD31 quantification, rat CD31/PECAM-1 antibody was used (Santa Cruz-Biotechnology; dilution: 1:50), whereas rabbit IBA1/AIF-1 (E4O4W) XP® (Cell Signaling Technology; dilution 1:1000) was used for the IBA1 detection. AQP4 was stained with rabbit AQP4 (D1F8E) XP® antibody (Cell Signaling Technology; dilution 1:200) and NeuN marker was determined by rabbit NEUN (D4G4O) XP® antibody (Cell Signaling Technology; dilution 1:400). Alexa Fluor 647 F(ab)2 fragment of goat anti-rabbit IgG (H+L) 2mg/ml (Invitrogen; dilution 1:1000) were used as secondary antibodies for CD31, IBA1, AQP4 and NeuN. Images were acquired using a Leica TCS SP5 inverted laser-scanning microscope on a Leica DMI 6000 CS module (Leica Microsystems) equipped with a HCX Plane-Apochromat lambda blue 63× oil-immersion objective with NA 1.4. Quantification of images was done using LasX software (Leica).

### Analysis of Immune Response on Mice

In mouse sera, specific anti-Spike SARS-CoV-2 total IgG was determined by ELISA test to test the immunogenicity of vaccines and determine the effect of hydroxychloroquine on the antibody production. ELISA tests were performed to determine End point titer. High-binding half-well plates (Greiner) were used. To determine specific anti-Spike SARS-CoV-2 total IgG, recombinant His-Spike SARS-CoV-2 protein (prepared in-house as described elsewhere^48^) was used for coating of the ELISA plates. Recombinant proteins were coated in PBS buffer at a concentration of 50 ng of designed protein per well. The total volume of the coated solution was 50 μl/well. Coated plates were incubated overnight at 4°C. The next day, plates were washed with PBS+0,05 % Tween20 using an ELISA plate washer (Tecan). Next, plates were blocked for 1h at RT with ELISA diluent (PBS+3%FBS) solution. Afterwards, the plates were again washed. Then, serial dilution (factor of 10) of mouse sera was added to the plates, where each dilution presented a certain titer value. For mouse sera ELISA diluent was used. Mouse sera were incubated at 4°C overnight. The next day, plates were washed and afterwards goat anti-mouse IgG (H+L)-HRP antibodies (Jackson ImmunoResearch), diluted 1:3000 in ELISA diluent solution. Plates were incubated for 1h at RT. Next plates were washed. After the final wash, TMB substrate was added, and the reaction was stopped with the addition of acid solution (3M H_3_PO_4_). Absorbance (A450 nm, A620 nm) was measured by Synergy Mx microtiter plate reader (Biotek). Next End point titers (EPT) were calculated. The end point titer was determined as the dilution above the value of the cutoff.

From absorbance data of control animals (non-treated animals), the cutoff value was determined^49^ by the formula, where is the mean of absorbance samples, SD is the standard deviation, and is the standard deviation multiplier. For a confidence level of 95% value 2,335 was used as the standard deviation multiplier.

### T-cell response on Mouse Splenocytes

To determine the antigen-specific cytotoxic CD8a^+^ T cells, spleens from immunized mice were harvested. Single cell suspensions from spleens without enzymatic treatment were obtained using the tissue dissociator gentleMACSTM Dissociator, according to the manufacturer’s instructions (MiltenyiBiotec, Bergisch Gladbach, Germany). CD8^+^ T cells from spleen cell suspension were isolated using the mouse CD8a^+^ T Cell Isolation Kit (Miltenyi Biotec; 130-104-075, Miltenyi Biotec, Bergisch Gladbach, Germany), according to the manufacturer’s instructions. Cells were isolated based on negative selection using LS columns, obtaining up to 10^8^ labeled cells. To determine SARS-CoV-2 Spike-specific cytotoxicity, mouse NIH-3T3 cells were seeded into 24-well plates (10^5^/well); the next day, the cells were transfected with pCG1-hACE2 (900 ng/well) and pCMV-TMPRSS2 (30 ng/well) plasmids. The following day, cells were infected with spike pseudovirus with a bioluminescent reporter (prepared in-house as described elsewhere^55^). The day after, isolated CD8a ^+^ T cells (10^5^/well) in RPMI1640 cell medium were added. After 24 h, bioluminescence was determined using IVISIII (Perkin Elmer, Waltham, MA, USA) after the addition of D-luciferin (500 μM), showing the state of the Spike-specific cytotoxicity of the CD8^+^ T cells isolated from vaccinated animals. Bioluminescence values are presented as average radiance (p/s/cm2/sr), which were determined using Living Image® software. From the average radiance values (ARV), the percentage of infected NIH-3T3-specific lysis was calculated using the following formula: % specific lysis =100 x (spontaneous death ARV-test ARV)/(spontaneous death ARV-maximal killing ARV). From the supernatants, mouse IL2 and mouse IFNγ levels were determined.

### Viral strains

SARS-CoV-2 EU1 strain (EPI_ISL_582134) was isolated from a nasopharyngeal swab of a patient during the first COVID-19 wave in March 2020. The virus was further propagated in Vero E6 cells (ATCC-CRL-1586) cultured in DMEM supplemented with 10% fetal calf serum (FCS), 1% penicillin/streptomycin, 200 mmol/L L-glutamine, 1% MEM-Non-Essential Amino Acids and 1% sodium pyruvate (all from Gibco, Thermo Fisher). A plaque assay was applied to determine the virus titer in plaque-forming units (PFU).

### Infection neutralization assay

Vero E6 cells were seeded at 1.5 × 10^4^ cells/well in a 96-well plate in supplemented DMEM medium and incubated overnight at 37 °C and 5% CO_2_. Serum samples were diluted 1:25 in culture medium, followed by a 5-fold serial dilution. Serum dilutions were mixed with SARS-CoV-2 virus to reach a multiplicity of infection (MOI) of 0.03 (450 PFU/15,000 cells/well) and incubated at 37 °C for one hour to enable virus neutralization. The inoculum was then incubated on Vero E6 cells for 1 h at 37 °C. Afterwards, the sample/virus mix was replaced by medium, and cells were cultured for 24 hours. As a positive control, cells were infected with the same MOI of the virus without incubating with serum samples, whereas uninfected Vero E6 cells represent mock. To stop the infection, the cells were rinsed once with PBS and fixed with 4% paraformaldehyde (ChemCruz) at RT for 15 min. After an additional PBS wash, fixed Vero E6 cells were permeabilized for 15 minutes at room temperature with 0.5% saponin (Roth) in PBS and blocked with PBS with 0.1% saponin and 10% BSA (Roth) overnight at 4 °C. The next day, VeroE6 cells were incubated for 2 hours at room temperature with a 1:1500 dilution of SARS-CoV-2 nucleocapsid antibody T62 (Sino Biological, Cat. No. 40143-T62) in PBS supplemented with 1% FCS. Cells were rinsed with wash buffer (PBS supplemented with 0.05% Tween-20 (Roth)) and incubated for one hour at room temperature with goat anti-rabbit IgG antibody, HRP conjugate (Merck KGaA, Cat. No. 12-348), 1:4000 diluted in PBS / 1% FCS. After thorough washing, 100µl TMB substrate (Invitrogen) was applied to each well and incubated at RT for 8 min in the dark. The reaction was stopped by adding 50 µl of 2 N H_2_SO_4_ (Roth), and colorimetric detection was performed on an Infinite F200 multi-plate reader (Tecan Group AG) at 450 and 560 nm. The data were fitted to a log(inhibitor) vs. response model with a variable slope in Prism v 8 (GraphPad).

### Statistical analyses

Data are presented as means ± SEM. One-way ANOVA followed by Tukey’s multiple comparisons test was used for the statistical comparison of data, if not otherwise stated. *P< 0.05, **P< 0.01, ***P< 0.001 ****P< 0.0001. Graphs were prepared using GraphPad Prism8.

## Acknowledgements

We thank K. Podgoršek for animal facility support and C.-C. Cheng for neutralization assay setup. Funding: Slovenian Research Agency (P4-0176, J4-4563), EU Horizon Teaming (101059842), EU H2020 FET Open VIROFIGHT (899619 to U.P. and R.J.), and BMBF C-NATM (03ZU1201BA to U.P.). Figures created with BioRender.com.

## Author contributions

D.L., Š.M. and R.J. conceived the study; D.L., Š.M., J.B., T.S. and A.G.U. performed most experiments; D.L., A.G.U. and S.O. conducted animal work; H.E. and J.P.Ž. performed cytotoxicity; V.F. performed IHC; M.B. led cytometry; R.B. performed neutralization assays; U.P. provided BNT162b2; R.J. supervised. All authors analyzed data and edited the manuscript.

## Competing interests

R.J., D.L., Š.M. and A.G.U. are inventors on a patent application related to this work; the authors declare no other competing interests.

## Reproducibility

Independent repeats specified in legends; key findings reproduced across donors and cohorts.

## Data availability

All data supporting the findings are available within the paper and **Extended Data**. RNA-seq data are deposited in GEO under accession **GSE277440**. Raw flow cytometry FCS files and source data for graphs are available from the corresponding author upon reasonable request.

## Code availability

Custom analysis used standard tools (Cutadapt, Kallisto, DESeq2, EnhancedVolcano, WebGestalt) with parameters described in the Reporting Summary; analysis scripts are available upon reasonable request.

## Supplementary Information

Supplementary Information includes plot of flow cytometry gating strategy and Table of antibodies used for flow cytometry in this study.

## Extended Data

**Extended Data Fig. 1:**
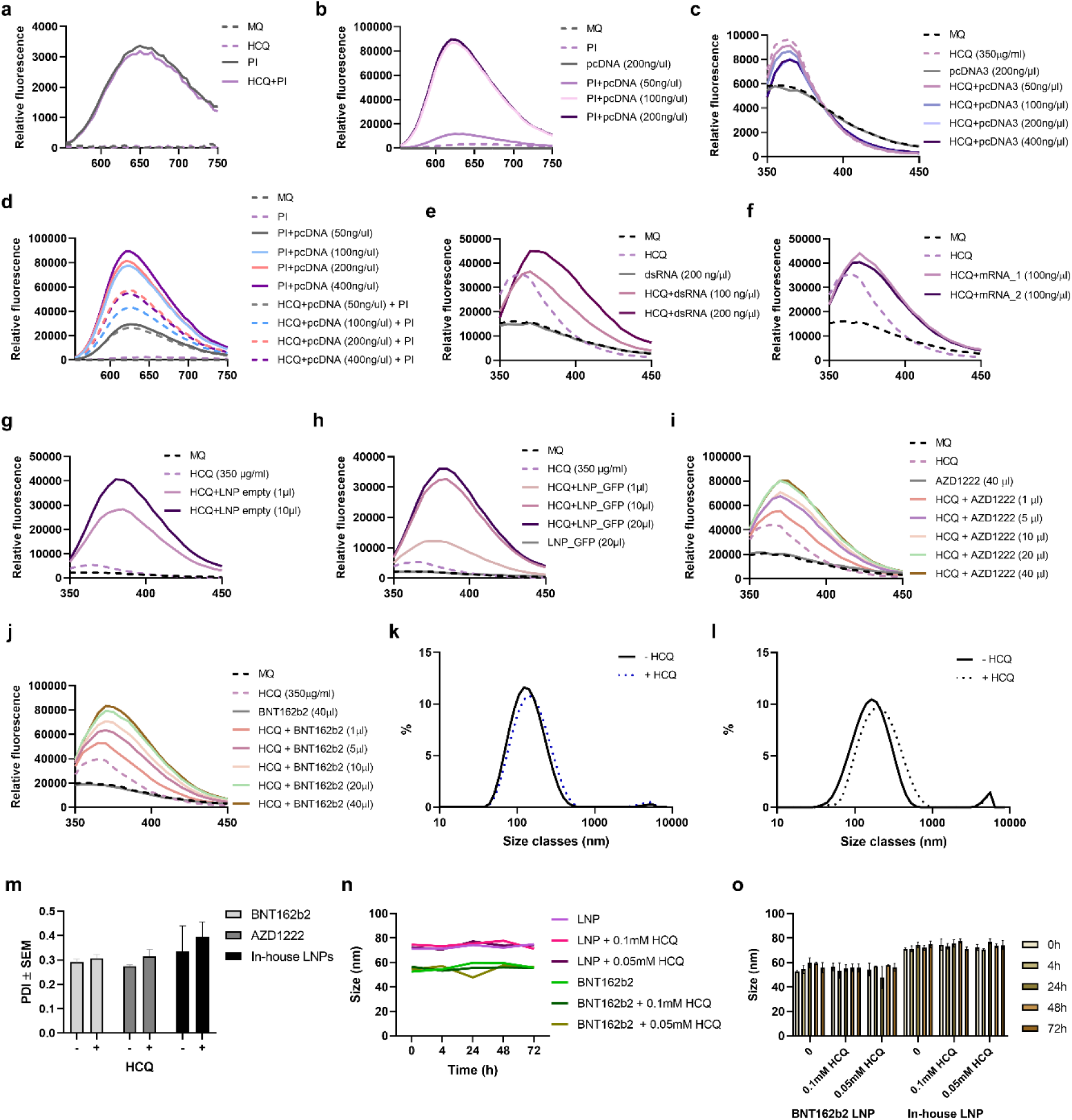
HCQ interacts with nucleic acids, viruses and LNPs without affecting their structural integrity. **a,** Emission spectra of PI (50 μM), MQ, and HCQ (1mM) upon excitation at 535 nm. **b,** Emission fluorescence spectra of the addition of plasmid DNA to PI (50 μM). **c**, Quenching of fluorescence intensity of HCQ (1 mM) by titration with plasmid DNA (pcDNA). **d,** Fluorescence emission spectra of PI (50 μM) in a mixture with plasmid DNA, both in the presence and absence of HCQ (1mM). **e,** Emission spectra of HCQ (1mM) upon the addition of dsRNA encoding for GFP. **f,** Emission spectra of HCQ (1mM) upon the addition of mRNA encoding for FLuc. **g,** Emission spectra of HCQ (1mM) after the addition of a gradual amount of empty LNPs. **h**, Emission spectra of HCQ (1mM) after the addition of a gradual amount of eGFP mRNA-LNPs. **i,** Fluorescent intensity of HCQ (1mM) after the addition of a gradual amount of AZD1222 viral vaccine. **j**, Emission spectrum of HCQ (1mM) after the addition of a gradual amount of BNT162b2 LNP mRNA vaccine. **k,** Intensity distribution of particles’ size classes for AZD1222. **l,** Intensity distribution of particles’ size classes for BNT162b2. **m,** Polydispersity indices (PDI) in the presence of HCQ. **n, o,** DLS determination of the size of LNP at different time-points in the presence of HCQ, indicating stable formulation.

**Extended Data Fig. 2:**
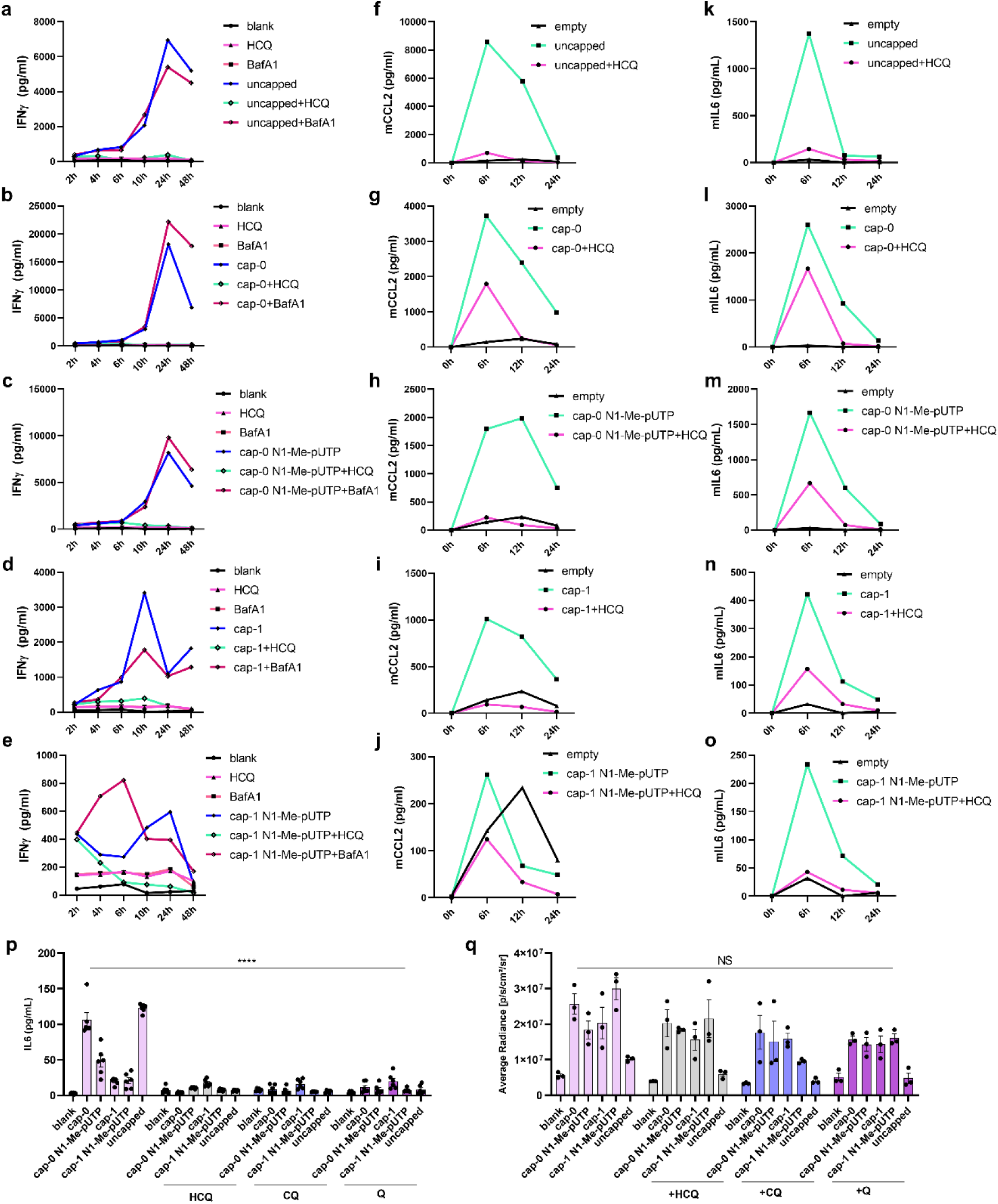
HCQ reduces inflammation, caused by different variants of mRNA in human cells. **a-e,** IFNγ of human PBMC (1μg of fLUC mRNA electroporation; n=3 donors; mean) in presence of HCQ (50μM) or bafilomycin A1 (100nM). **f-j,** Mouse CCL2 in mice sera (5μg of LNPs, containing different variants of fLUC mRNA+HCQ (60 mg/kg of body weight) n=3 mice/group; mean). **k-o,** Mouse IL6 in mice sera (5μg of LNPs, containing different variants of fLUC mRNA+HCQ (60 mg/kg of body weight) n=3 mice/group; mean). **p**, IFNγ of human PBMC (1μg of fLUC mRNA electroporation; n=3 donors; mean±SEM) in the presence of HCQ, CQ (chloroquine) or Q (quinacrine) (50μM). **q**, Bioluminescence measurement 24 hours later, PBMC electroporation (n=3 donors; mean±SEM).

**Extended Data Fig. 3:**
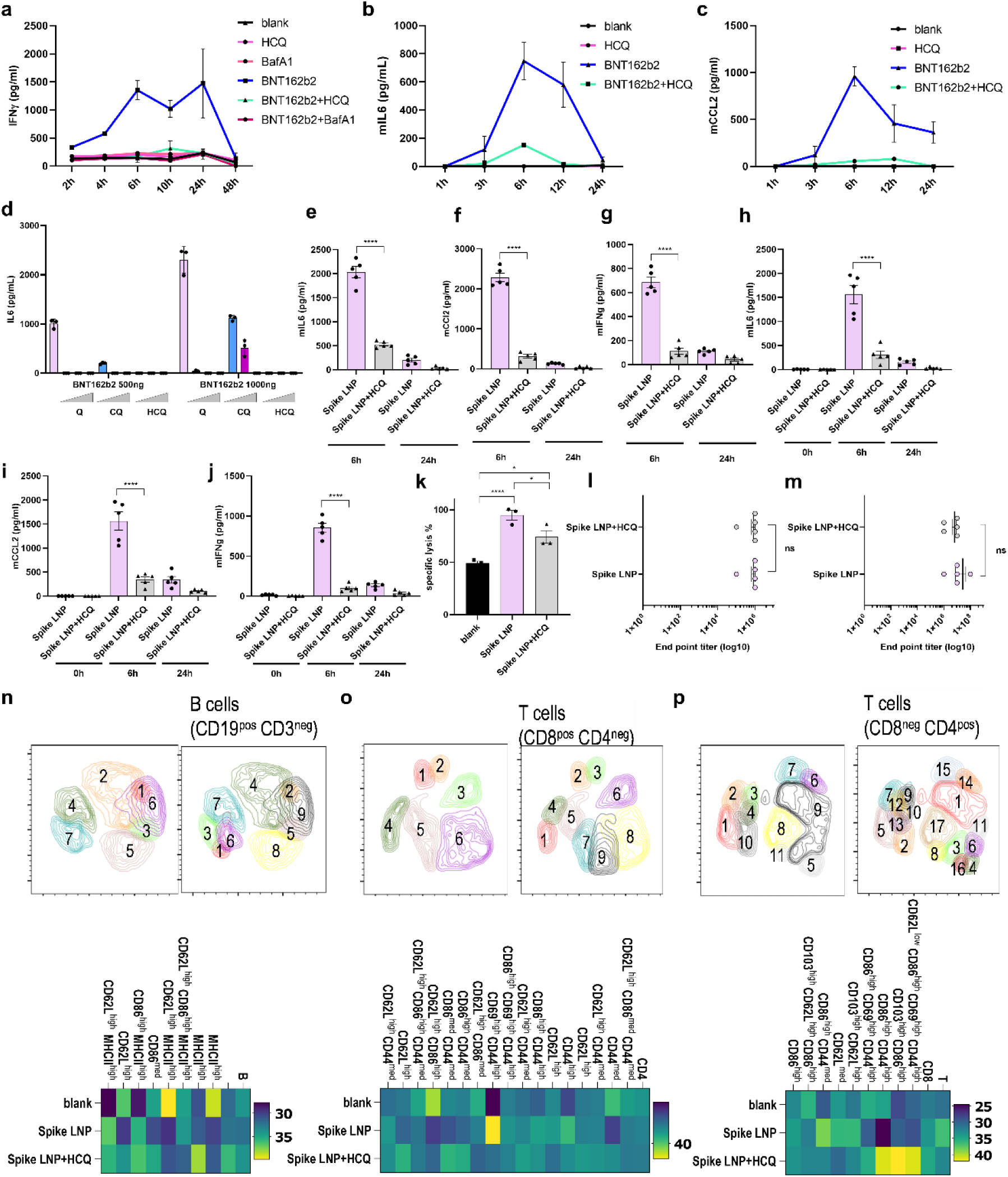
HCQ reduces mRNA vaccine-induced inflammation. **a,** Human IFNγ of PBMCs (mean±SEM; n=2 donors), stimulated with BNT162b2 vaccine (500ng) with the addition of HCQ (50μM) or bafilomycin A1 (100nM). **b-c**, Mouse IL6 (**b**) and mouse CCL2 (**c**) injected with mixture of BNT162b2 (4µg) and HCQ (60 mg/kg of body weight) (mean±SEM; n=3mice). **d**, Human IL6 of PBMCs (mean±SEM; n=2 donors), stimulated with BNT162b2 vaccine with the addition (1μg/ml, 5 μg/ml an 10 μg/ml) of HCQ, CQ or Q. **e-g**, Cap-0 unmodified Spike mRNA/LNP (4 µg) ± HCQ (60 mg/kg of body weight) intramuscularly (mean±SEM; n=5 mice/group); mIL6 (**e**), mCCL2 (**f**), mIFNγ (**g**) at 6 h/24 h after prime. **h-j**, mIL6 (**h**), mCCL2 (**i**), mIFNγ (**j**) at 0 h/6 h/24 h after boost. **k**, CD8⁺ cytotoxicity against Spike-pseudovirus–infected cells (day 42). **l–m**, End-point anti-Spike IgG titres after prime (**l**) and boost (**m**). **n-p**, Spectral flow cytometry UMAP of CD3 cells from spleens of Spike LNP vaccinated animals; CD19 cells (**n**) CD8^+^ T cells (**o**) or CD4^+^ T cells (**p**).

**Extended Data Fig. 4:**
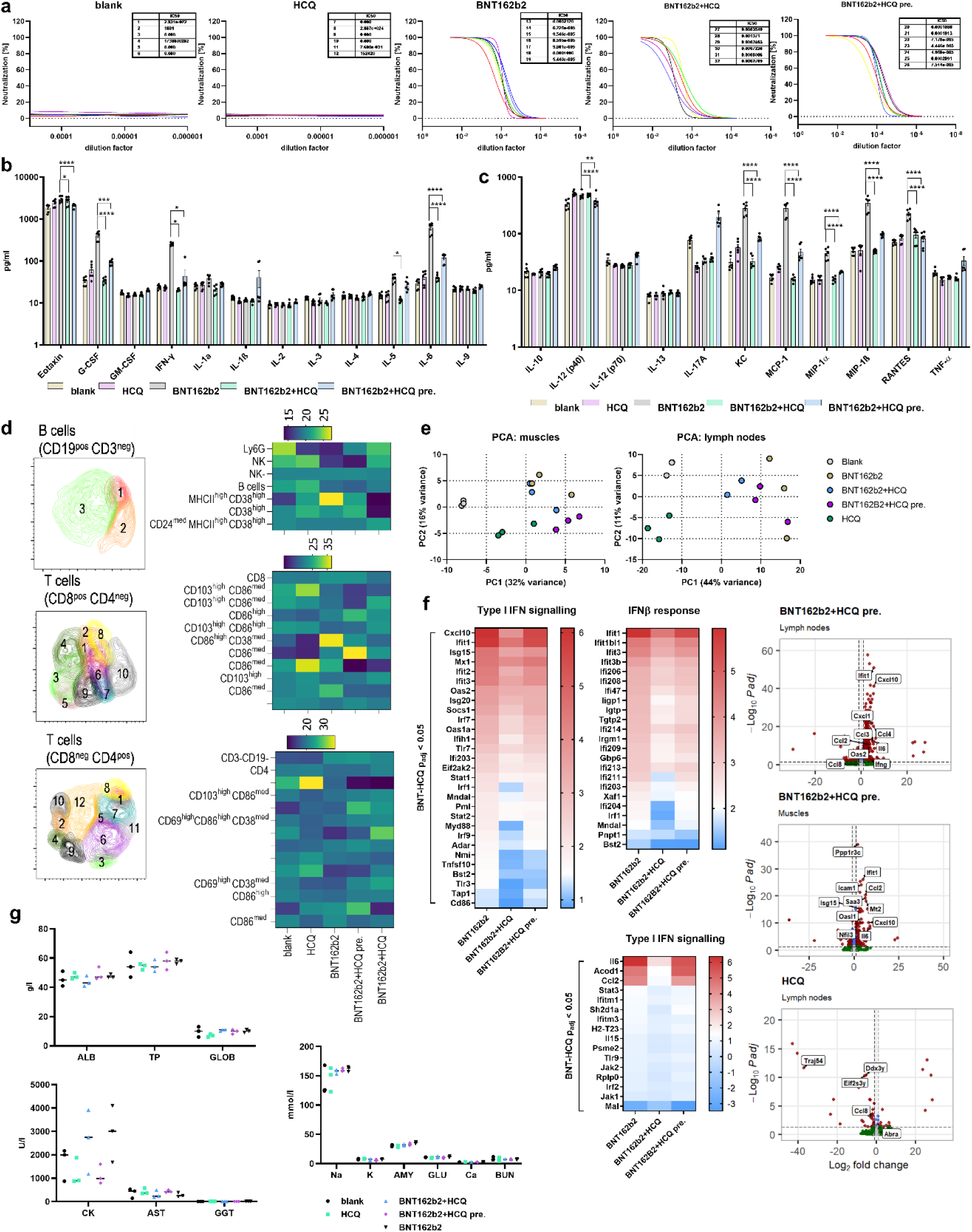
HCQ does not alter BNT162b2 vaccine efficiency. **a,** Wuhan strain SARS-CoV-2 virus neutralization by immunized mice sera. **b-c,** BNT162b2 intramuscular immunization (4μg mRNA BNT162b2 vaccine and HCQ (60mg/kg body weight) intramuscularly (mena±SEM; n=6 mice). **d**, Spectral flow cytometry analysis of spleen cells, obtained from BNT162b2 vaccinated animals (n=6). **e,** PCA analysis for lymph nodes and muscles. **f**, Enhanced volcano plots for BNT162b2+HCQ pre. and HCQ only groups, Type I IFN signaling heatmap for lymph nodes and IFNβ response for lymph nodes. g, Biochemical blood values from immunized animals BNT162b2 (4μg/animal) with or without HCQ (60 mg/kg of body mass; mean±SEM; n=6). ALB, albumins; TP, total proteins; GLOB, globulins; ALP, alkaline phosphatase; ALT, alanine aminotransferase; GGT, gamma glutamyl transferase; AMY, amylase; TBIL, total bilirubin; CRE, creatinine kinase; BUN, blood urea nitrogen; Glu, glucose.

**Extended Data Fig. 5.**
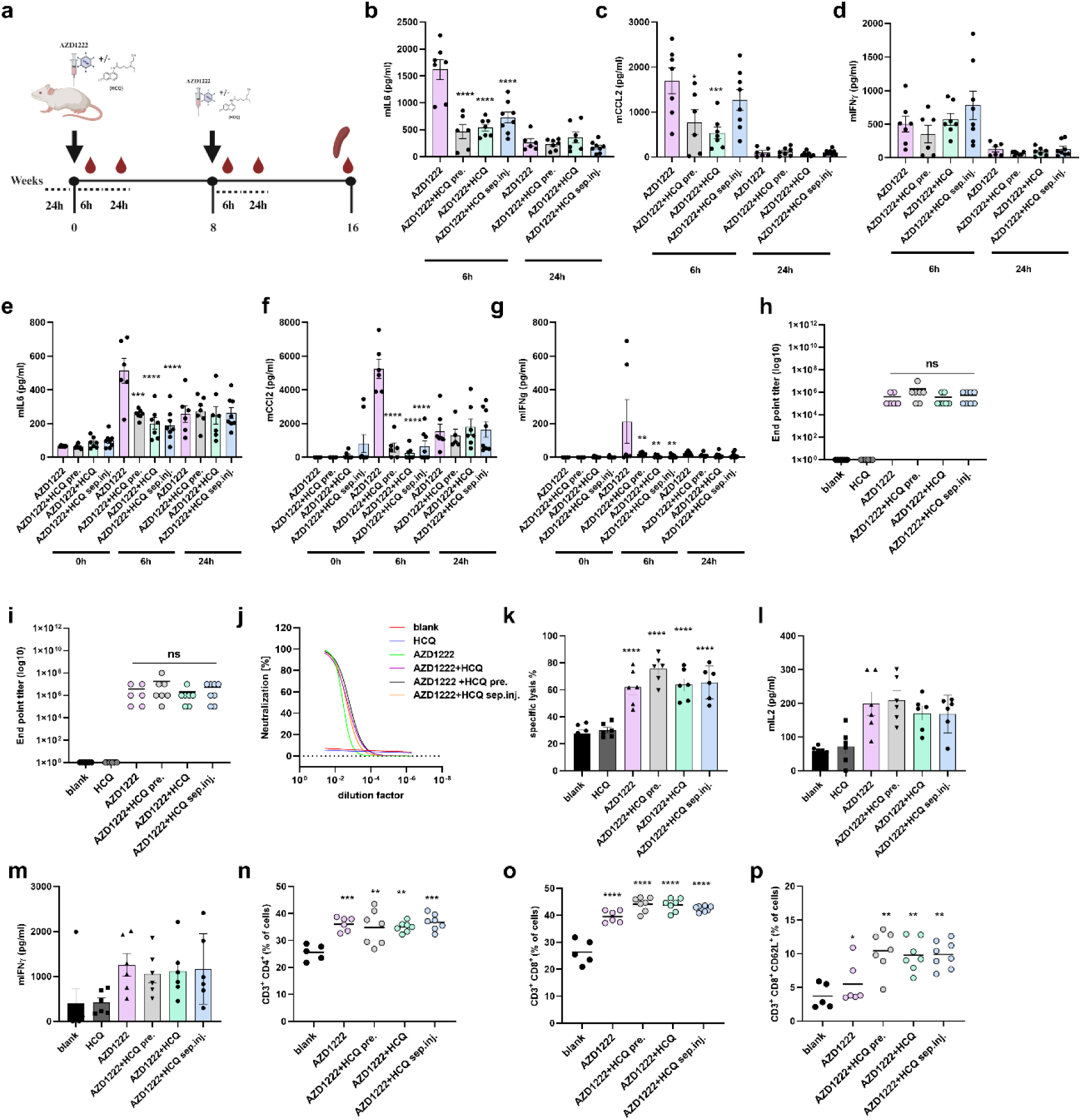
HCQ suppresses adenoviral vaccine reactogenicity while maintaining efficacy. **a**, AZD1222 prime–boost with HCQ co-formulated, contralateral (sep. inj.) or prophylactic (HCQ pre). **b–d**, Cytokines after prime; **e–g**, after boost (8 weeks). **h–i**, Anti-Spike IgG titres (prime, boost). **j**, Live-virus neutralization. **k–m**, Splenocyte cytotoxicity and cytokines. **n–p**, Flow cytometry (CD4⁺, CD8⁺, CD62L⁺CD8⁺) at week 16. n=6; mean±SEM.

**Extended Data Fig. 6:**
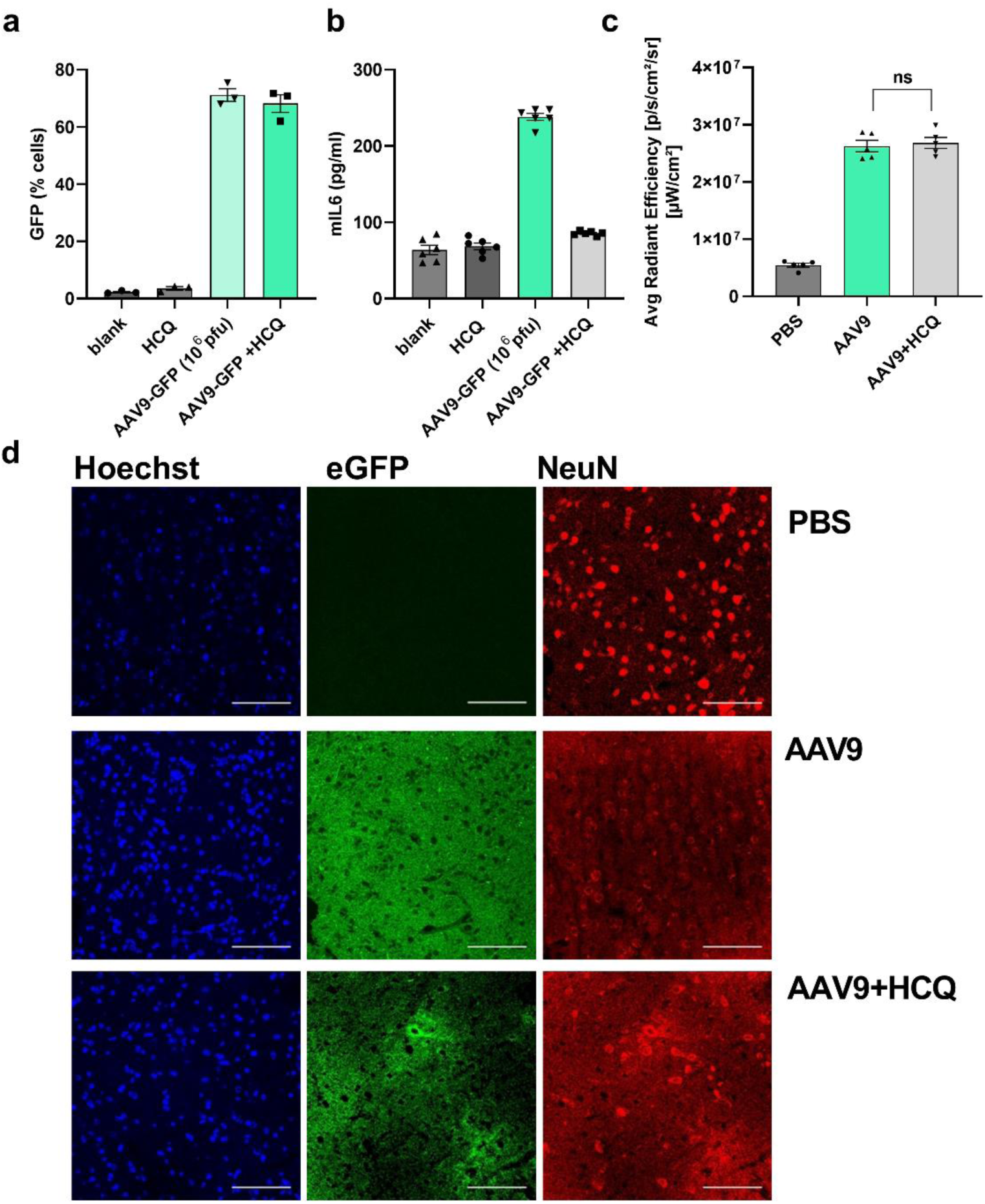
HCQ mitigates AAV9 induced damage. **a,** Flow-cytometric determination of eGFP expression in mouse N2A cells (AAV9-eGFP; 10^6^pfu+ HCQ (50μM); mean±SEM; n=3). **b**, Mouse IL6 values of N2A cells (AAV9-eGFP; 10^6^pfu+ HCQ (50μM); mean±SEM; n=6). **c**, In vivo fluorescence measurement of eGFP fluorescence signal in vivo 30 days post AAV9-eGFP injection (1×10¹⁰ pfu, cortex; HCQ (60mg/kg body weight) (mean±SEM; n=5). **d**, NeuN neuronal marker staining, determining the level of neuronal density. Scale bars correspond to 100 μm. Representative images are shown.

**Extended Data Fig. 7:**
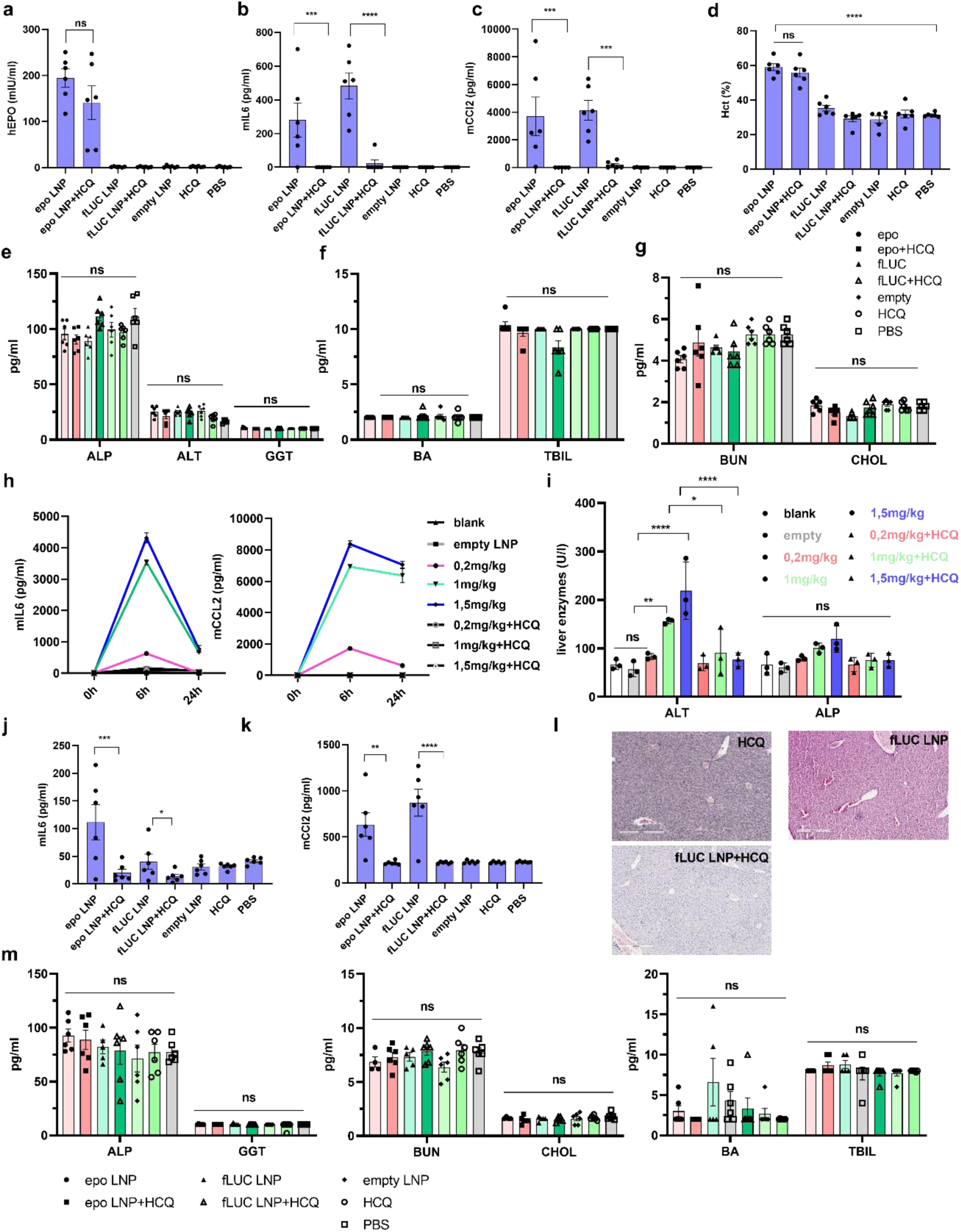
The effect of erythropoietin (epo) or fLUC mRNA delivery via LNPs with simultaneous HCQ administration in vivo. **a-d,** Epo or fLUC coding cap-1 unmodified mRNA LNP (0,2mg/kg of body weight of) + HCQ (60mg/kg body weight) every 3 days for 1 week in mice intravenously (mean±SEM; n=6 mice). Human Epo (**a**), mouse IL6 and (**b**) mouse CCl2 (**c**) 6 h later after the second LNP injection. Hematocrit 8 days following the LNP injection (**d**). **e-g,** Blood liver profile 3 days after the LNP injection. **h-i**, Different doses of eGFP mRNA LNP (cap-1 unmodified) + HCQ (60mg/kg body weight) intravenously (mean±SEM; n=3 mice). Mouse IL6 and mouse CCl2 at different timepoints (**h**). **i**, ALT and ALP 2 days following the LNP injection (**i**). **j-m**, Epo or fLUC coding cap-1 unmodified mRNA LNP (1mg/kg of body weight of) + HCQ (60mg/kg body weight) every 3 days for 1 week in mice intravenously (mean±SEM; n=6 mice). Mouse IL6 and (**j**) mouse CCl2 (**k**) 24h after the first LNP injection. Liver histology 8 days post-injection. Representative images are shown. Scale bar corresponds to 300 µm (**l**). Blood liver profile 3 days after the second LNP injection (**m**); ALP-alkaline phosphatase; ALT-alanine aminotransferase; GGT-gamma-glutamyl transferase; BA-bile acids; TBIL-total bilirubin; BUN-blood urea nitrogen; CHOL-cholesterol.

**Extended Data Fig. 8:**
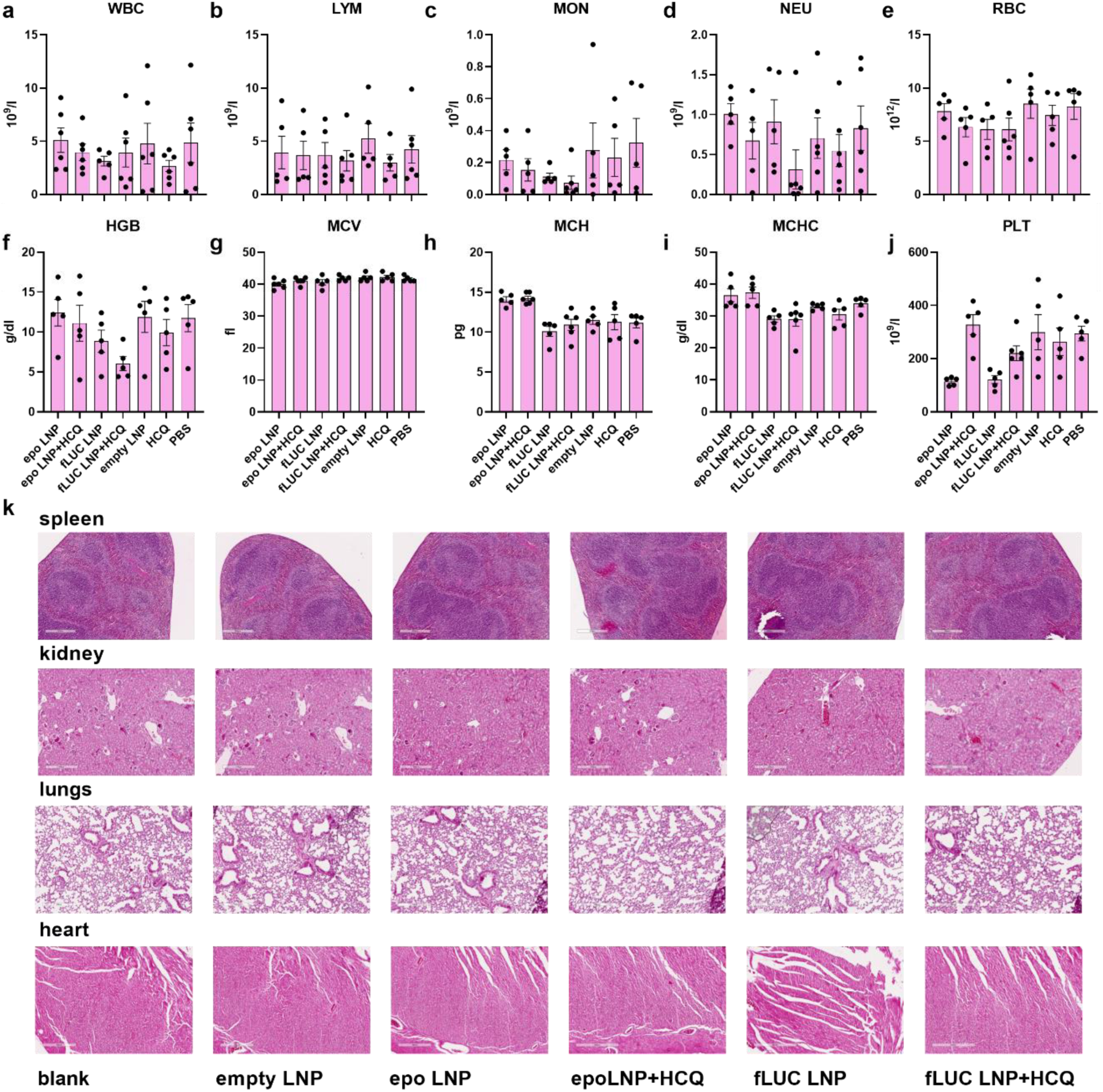
HCQ administration in vivo lowers the side effects of mRNA LNP therapy. **a-k,** Epo or fLUC coding cap-1 unmodified mRNA LNP (1mg/kg of body weight of) + HCQ (60mg/kg body weight) every 3 days for 1 week in mice intravenously. Blood count 8 days post-LNP injection (mean±SEM; n=6 mice). (WBC-White Blood Cell Count; LYM-Lymphocytes; MON-Monocytes; NEU-Neutrophils; RBC-Red Blood Cell Count; HGB-Hemoglobin; MCV-Mean Corpuscular Volume; MCH-Mean Corpuscular Hemoglobin; MCHC-Mean Corpuscular Hemoglobin Concentration; PLT-Platelets) (**a-j**). Tissue H&E staining 8 days post-LNP injection. Representative images are shown. Scale bar∼300 µm (**k**).

**Extended Data Fig. 9:**
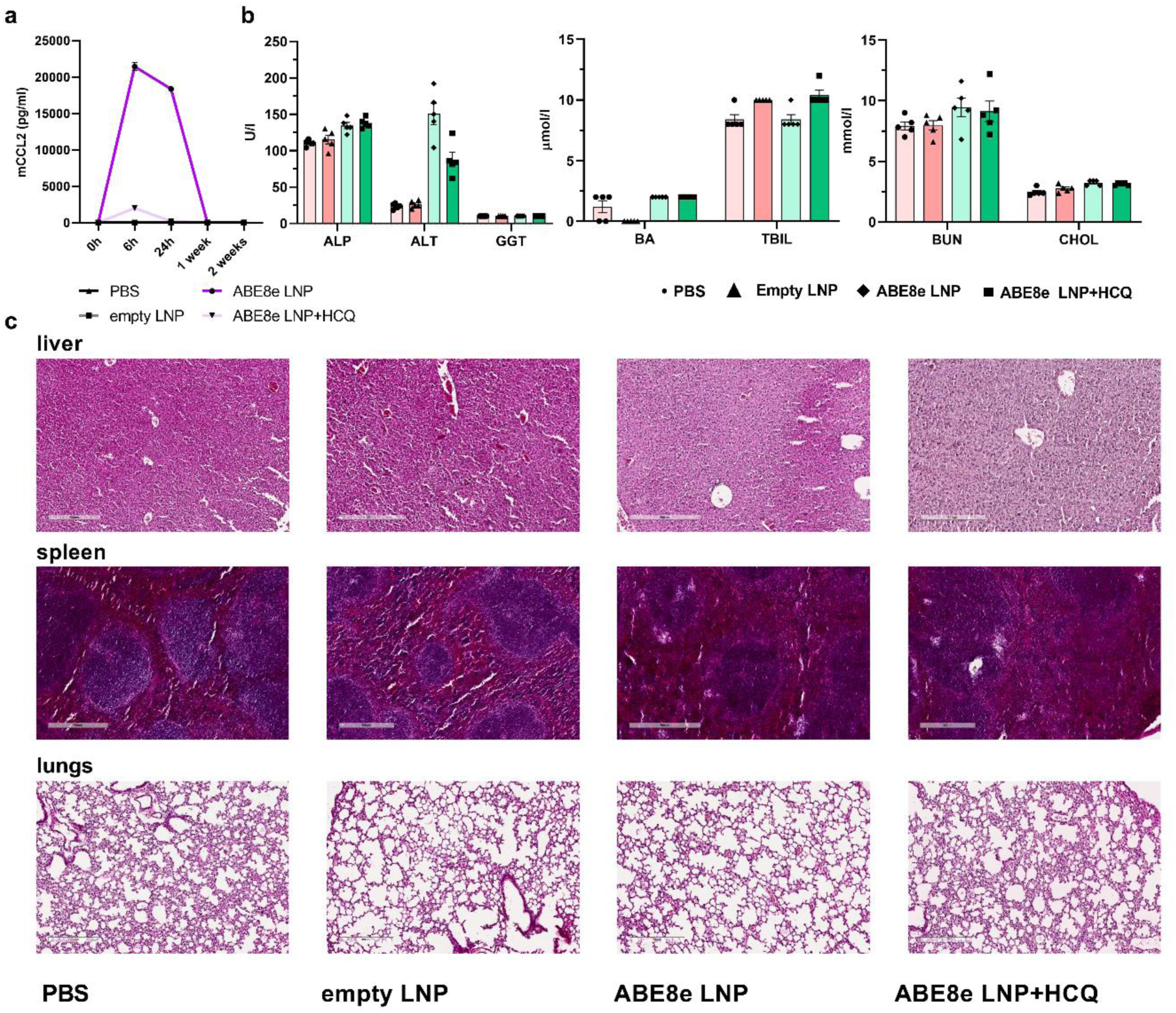
Effect of HCQ in genome editor delivery. **a-c,** ABE8e + *Pcsk9* gRNA mRNA/LNP (1.5 mg/kg of body weight) ± HCQ (60 mg/kg body weight). Re-dosage was performed three days later (mean±SEM; n=5 mice). Mouse CCl2 (**a**). Liver biochemical profile at day 4. (ALP-alkaline phosphatase; ALT-alanine aminotransferase; GGT-gamma-glutamyl transferase; BA-bile acids; TBIL-total bilirubin; BUN-blood urea nitrogen; CHOL-cholesterol) (**b**). Tissue H&E staining 21 days post LNP injection. Representative images are shown from the end of the experiment at 21 days. Scale bar∼300 µm (c).

## Notes

### Competing Interest Statement

R.J., D.L., S.M. and A.G.U. are inventors on a patent application related to this work; the authors declare no other competing interests.

